# Local ionic conditions modulate the aggregation propensity and influence the structural polymorphism of alpha-synuclein

**DOI:** 10.1101/2024.11.03.621709

**Authors:** Maria Zacharopoulou, Neeleema Seetaloo, James Ross, Amberley D. Stephens, Giuliana Fusco, Thomas M. McCoy, Wenyue Dai, Ioanna Mela, Ana Fernandez Villegas, Anne Martel, Alexander F. Routh, Alfonso De Simone, Jonathan J. Phillips, Gabriele S. Kaminski Schierle

**Affiliations:** Department of Chemical Engineering and Biotechnology, University of Cambridge, Philippa Fawcett Drive, Cambridge, UK; Living Systems Institute, University of Exeter, Stocker Road, Exeter, UK; School of Molecular and Cellular Biology and Astbury Centre for Structural Molecular Biology; Department of Chemistry, University of Cambridge, Cambridge CB2 1EW, UK; Department of Life Sciences, Imperial College London, London SW7 2AZ, UK; Institut Laue Langevin, Grenoble, France

**Author notes:** Corresponding Author Gabriele Kaminski Schierle.

**Keywords:** alpha-synuclein, neurodegeneration, protein aggregation, fibrillation, polymorphism, intrinsically disordered proteins, conformational dynamics, water mobility, solvation

## Abstract

Parkinson’s Disease (PD) is characterized by the aggregation of alpha-synuclein (aSyn), a presynaptic protein that transitions from a disordered monomer into beta-sheet rich amyloid fibrils. The precise triggers and mechanisms underlying aSyn misfolding and aggregation remain unclear, hindering the development of effective therapeutics. Monomeric aSyn is an intrinsically disordered protein (IDP) with high conformational flexibility. Local environmental factors, such as ion concentrations, can influence the conformational ensemble of aSyn, impacting its aggregation propensity and resulting in fibril polymorphism. In this study, we explore the impact of physiologically relevant ions, mainly Ca^2+^ and Na^+^, on the aggregation kinetics, monomer structural dynamics, and fibril polymorphism of aSyn. Using ThT fluorescence assays, we demonstrate that all ions accelerate aSyn aggregation, with Ca^2+^ having the most significant effect. Using Heteronuclear Single Quantum Correlation Nuclear Magnetic Resonance (^1^H-^15^NHSQC NMR) spectroscopy, we validate the specific binding of Ca^2+^ ions at the C-terminus, whereas Na^+^ ions display non-specific interactions along the sequence of aSyn. Small-angle neutron scattering (SANS) and hydrogen-deuterium exchange mass spectrometry (HDX-MS) further reveal that Na^+^ and Ca^2+^ induce distinct conformational changes in the aSyn monomer, with Na^+^ leading to more extended structures and Ca^2+^ promoting a moderate extension of the protein. Molecular dynamics simulations (MD) corroborate these findings, showing that Na^+^ ions increase the protein’s extension, particularly between the non-amyloid beta component (NAC) region and the C-terminus, whereas Ca^2+^ ions bias the ensemble towards a more moderately elongated structure. Using MD, we further investigate the local environment and in particular the solvent effect and show the water persistence times in the hydration shell are also increased in the presence of Ca^2+^ ions, indicating that the aggregation propensity of the monomer is due to a combination of conformational bias of the monomer and solvent mobility. Atomic force microscopy (AFM) of aSyn fibrils formed under these different ionic conditions reveal distinct fibril polymorphs, suggesting that ion-induced conformational biases in the monomer contribute to the diversity of fibril structures. Collectively, these findings underscore the pivotal influence of the local ionic milieu in shaping the structure and aggregation propensity of aSyn, thus offering valuable insights into the molecular underpinnings of PD and potential therapeutic avenues aimed at manipulating aSyn conformational dynamics.

## INTRODUCTION

Current treatments for Parkinson’s Disease (PD) patients are symptomatic, as the detailed molecular mechanism leading to PD pathology remains elusive ^1,2^. Evidence points towards the aggregation of a small presynaptic protein, alpha-synuclein (aSyn), as it transitions from its functionally disordered form into beta-sheet rich amyloid fibrils which are present in Lewy body (LB) inclusions, one of the hallmarks of PD ^3–5 6–14^. However, many unanswered questions remain with regards to aSyn aggregation in neurons and, in particular, when the soluble, functional form of monomeric aSyn starts to misfold into a structure that consequently facilitates aSyn aggregation. Elucidating the triggers for aSyn misfolding and the associated monomeric structures and dynamics will aid the design of effective therapeutics.

Monomeric aSyn is an intrinsically disordered 14.4 kDa protein, consisting of 140 amino acid residues which can be divided into three specific categories: a highly positively charged amphipathic N-terminus (1-60), a central hydrophobic core (61-95) known as the non-amyloid beta component (NAC) region, and an acidic, C-terminal tail (96-140) ^15,16^. aSyn has remarkable conformational flexibility, structural plasticity and no unique structure, in contrast to well-folded proteins ^17^. Due to this conformational plasticity, an ensemble of different protein structures is expected to co-exist, dependent on the protein’s local environment and binding partners ^18,19^. The inherent fluctuations in the structure of an unfolded protein permit residues that are separated in the primary sequence to encounter one another in space. There is increasing evidence that residual structure and intramolecular interactions exist within monomeric aSyn, despite its conformational flexibility ^20–23^. These interactions are stabilized by hydrogen bonds, electrostatic and hydrophobic interactions and account for aSyn’s smaller radius of gyration, compared to the predicted random-coil 140-residue protein, suggesting a partially folded structure ^24^. The partial folding of the protein has been proposed to modulate the fibrillation propensity of aSyn ^25^. It is thus important to understand whether and how a bias in the conformational ensemble of aSyn, modulated by an alteration of these long-range interactions, may impact the aggregation of aSyn and the resulting fibrils.

Indeed, even though the trigger(s) that initiate the misfolding of soluble disordered aSyn into insoluble fibrils are still unknown, there is growing evidence that pathology is initiated by the disruption of the monomeric aSyn conformation ^26,27^. The conformational ensemble of aSyn is further expected to be skewed towards different conformers in different cellular compartments, where diverse microenvironments are maintained (different concentrations of ions, pH, or binding partners) ^28^. The interaction of the protein with the surrounding solvent is also pertinent, as ions influence the mobility of water molecules in the solvation shell of a protein. This interaction is especially relevant for intrinsically disordered proteins such as aSyn, which exhibit significantly larger solvent-accessible areas in comparison to globular proteins of similar size ^29^.

There are several possible routes for aSyn to encounter different environmental conditions. The release of aSyn into the extracellular space through routes such as cell death and release of cellular contents, exocytosis, or exosome release, may lead to aSyn being exposed to high salt and high calcium concentrations ^30,31^. Subsequent uptake by endocytosis into the endosomal/lysosomal pathway would expose aSyn to a low pH environment ^32^. Furthermore, calcium dysfunction^33^ and mitochondrial dysfunction^34^ are hallmarks of aging cells ^35–37^and may lead to alterations in the cellular environment and thus lead to aSyn structures that are trapped in a more aggregation-prone conformation.

Besides its impact on the monomer conformation, the local environment has also been shown to affect fibril polymorphism. Structural studies *in vitro* have demonstrated that the presence of salt ions can yield distinct aggregate structures, with twisted fibrils forming in the presence of salt and ribbon-like fibrils forming in the absence of salt, and thereby exhibiting different levels of toxicity ^38–40^. The formation of these fibril polymorphs has also been observed and reported in the recent publications of high-resolution structures using cryoEM, where different fibril structures are formed in almost every study ^41–46^, driven by the distinct physicochemical conditions in which the fibrils are grown (e.g. salt concentration, crystallization factors). Fibril polymorphism can be attributed either at the protofibril-level structure (kernel structure), or to how the protofibrils intertwine with each other to give rise to full amyloid fibrils ^44^. The existence of different kernels at the protofibril level suggests that distinct fibril polymorphs arise not only from differences in the way the two protofibrils twist around each other, but also from structural characteristics that precede protofibril association. Thus, the distinct polymorphs may stem from the structural characteristics of the oligomers and/or the original monomer conformation. Recently, the structure of *ex vivo* aSyn fibrils from post-mortem tissue of patients with Multiple Systems Atrophy (MSA), Dementia with Lewy Bodies (DLB), and PD has been resolved via cryo electron microscopy (EM) ^47–51^, providing further evidence that different fibril polymorphs are related to different disease phenotypes.

In light of the structural plasticity of the aSyn monomer and the diversity of its aggregation products, we aim to understand how the local environment affects the conformational ensemble of aSyn, the aggregation propensity of the protein, and the structure of the formed fibrils. In our previous publications, ^52,53^ we investigated the effect of the physiologically relevant Ca^2+^ ion binding on the aggregation propensity and conformational state of aSyn. By studying the aggregation kinetics of a panel of familial mutants, we uncovered distinct aggregation kinetics in response to the presence of Ca^2+^. We further established that a bias in the exposure of the aSyn monomer (especially at the N-terminus and NAC region) correlated positively with the protein’s aggregation propensity. We observed a correlation between the aSyn aggregation kinetics in distinct ionic environments, building up towards mimetics of the intracellular, extracellular, and lysosomal solvent environment^50^. Furthermore, we showed that the solvent dynamics were an important factor affecting aSyn aggregation: ions that decreased the water mobility in the solvation shell of aSyn led to an increase in its aggregation rate ^54^. Here, we focus the study of the conformational dynamics of aSyn for the physiologically relevant NaCl and KCl salts, to elucidate the added effect of these environmental parameters on the aggregation propensity of aSyn.

We study the aggregation kinetics of aSyn in the different ionic conditions, related to the intracellular and extracellular space. We investigate the monomer structural dynamics of aSyn in the same conditions by Hydrogen-Deuterium Mass Spectrometry (HDX-MS), Heteronuclear single quantum coherence Nuclear Magnetic Resonance (^1^H-^15^N HSQC NMR) and Small-angle Neutron Scattering (SANS) and relate the differences in conformation to the protein’s aggregation behaviour. Similar to our previous work^53^, recent advances in instrument development allowed us to collect HDX-MS data at high structural and temporal resolution, with HDX labelling in the millisecond regime coupled with soft fragmentation in the gas phase (electron-transfer dissociation -ETD)^55,56^. We further model the monomeric conformation of aSyn in the same environmental ion conditions, using coarse-grained MD simulations, and extract structural information of the conformational ensemble of aSyn, as well as its interactions with the ions and the solvent. Finally, at the fibril level, we probe the structural polymorphism of the aSyn fibrils formed in their distinct environmental conditions via Atomic Force Microscopy (AFM).

We conclude that the solution conditions assessed in this study (Na^+^, Ca^2+^, Na^+^ and Ca^2+^, K^+^) lead to distinct local conformational changes of the aSyn monomer that influence the aggregation kinetics and the polymorphism of the formed fibrils. We observe the aggregation rate of aSyn in CaCl^2^, NaCl, and KCl is increased in comparison to a “no salt” environment, with CaCl^2^ having a much stronger effect that NaCl and KCl. Both NaCl and KCl have the same effect on the aggregation kinetics, so we have here focused our efforts on elucidating the effect of Ca^2+^ and Na^+^ on aSyn monomer structure and dynamics. We observe that Na^+^ has no specific binding site, in contrast to Ca^2+^, which has its binding site at the C-terminus of aSyn retained in the presence of Na^+^. Both Na^+^ and Ca^2+^ ions induce an extension of the aSyn monomer structure: the addition of only Na^+^ induces the largest structural extension, followed by Na^+^ and Ca^2+^, and Ca^2+^ only. Furthermore, we show that the solvent mobility in the hydration shell of aSyn is reduced in the presence of Ca^2+^ compared to Na^+^. By correlating these observations on the monomer to the aggregation kinetics assays we suggest that the rate of aSyn aggregation is influenced by a combination of a structural extension of the aSyn monomer, as well as by the solvent mobility in the hydration shell of the protein. This further leads to the formation of distinct fibril polymorphs in the presence of different ions, with disease-relevant, more twisted fibrils formed in the presence of Ca^2+^. These findings suggest that the ionic milieu impacts not only the aggregation kinetics, but also the ultimate aggregation product. Our results therefore highlight the importance of the local environment on aSyn aggregation and suggest ways for therapeutic intervention by designing molecules that: (i) stabilise a more compact monomer conformation, (ii) regulate the ion household in the cell, (iii) and/or modulate the hydration shell around the protein (osmolytes) and thus increase the protein’s mobility directly to prevent aggregation.

## RESULTS

### The aggregation kinetics of aSyn is influenced by the local ionic milieu

We have previously investigated the aggregation and structure of aSyn monomer in complex ionic environments, which mimic different cellular compartments, showing increased aggregation at low pH and high Ca^2+^ concentrations ^52,53^. Here, we want to further probe the mechanisms of aggregation kinetics twinned with monomer structural assessment by studying a more simplistic environment that is easier to model by MD simulations. We first investigate how the major ion components of the local physiological environment, namely Na^+^, K^+^, and Ca^2+^ ions, influence the aggregation rate of aSyn, building up to *in vitro* ion mimetics of the intracellular and extracellular environment. To do so, we first employ a ThT fluorescence-based assay, widely used in the amyloid field ^57^. Briefly, the ThT molecule fluoresces when bound to rich fibrillar β-sheet structures, such as those found in amyloid fibrils, and thus the assay provides us with a tool to measure aSyn aggregation rates. The aggregation kinetics typically follow a sigmoidal curve. Aggregation kinetics are fitted, and three parameters are extracted, namely: t^lag^ (the time spent in the lag phase, corresponding to nucleation), k (the slope of the exponential phase, corresponding to fibril elongation), and t^50^ (the mid-point of the exponential phase, also corresponding to elongation). We probe the aSyn aggregation rates for five separate conditions: aSyn in Tris buffer only, pH 7.4, aSyn in 2 mM CaCl^2^, aSyn in 150 mM KCl (corresponding to the intracellular space, **Figure S2**), aSyn with 150 mM NaCl, and aSyn with 150 mM NaCl and 2 mM CaCl^2^ buffer solutions (corresponding to the extracellular space) (**Figure 1a-d**). We also measure the remaining monomer concentration at the end of the ThT-based assays as an orthogonal method to determine the extent of aSyn aggregation (**Figure 1e**). Distinct aggregation kinetics can be identified for each condition, indicating the different effects ions have on the aggregation propensity of aSyn.

**Figure 1:**
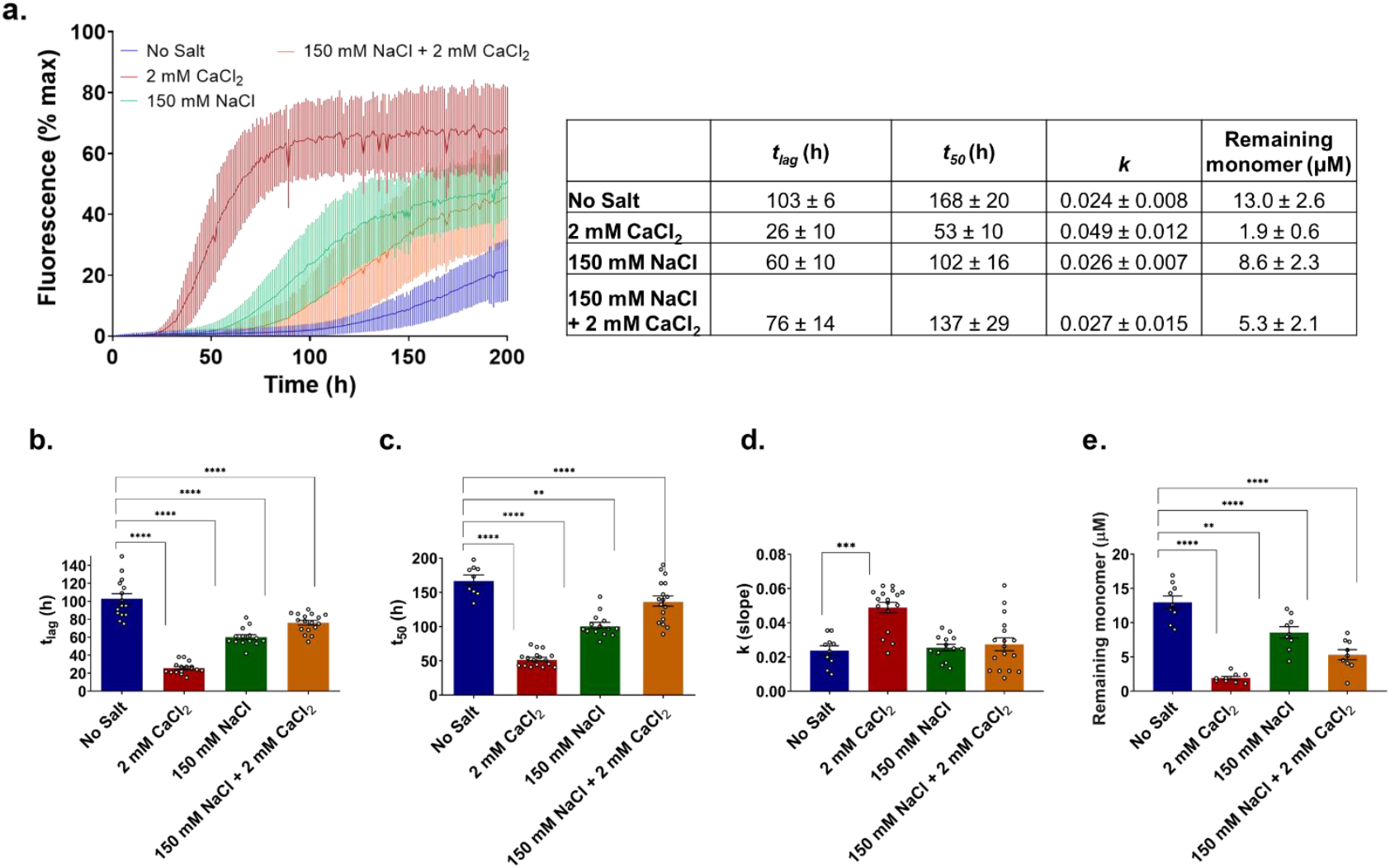
ThT aggregation kinetics of WT aSyn is increased in the presence of all ions (Na^+^, Ca^2+^), with Ca^2+^ displaying the highest increase. The aggregation kinetics of aSyn were determined by measuring ThT fluorescence intensity which was plotted as % of maximum fluorescence. 20 μM aSyn was incubated with 20 μM ThT in a 96 well plate with agitation at 300 rpm for 5 minutes before each read out every hour for 250 hours. The conditions studied are aSyn in 20 mM Tris pH 7.4, with addition of 2 mM CaCl_2_, 150 mM NaCl, 150 mM NaCl and 2 mM CaCl_2_. At least 6 replicates across three biological repeats were collected per condition. **a**. Kinetic traces. The average between traces of the same condition is shown in the graph and errors indicate 1 s.d., **b**. lag time (t_lag_), **c**. time to reach 50 % of maximum aggregation (t_50_) and **d**. the slope of the curve *k* (calculated by fitting Equation 1), and the mean plus error (1 s.d.) are displayed in the graphs. **e**. Remaining monomer concentration (μM) at the end of the aggregation assay was determined using SEC-HPLC, where 25 μL of sample from each well in the ThT assays was analysed using an AdvanceBio SEC 130Å column in 20 mM Tris pH 7.4 at 0.8 mL min^-1^. An ordinary ANOVA was used to calculate statistical significance between samples and significant differences are reported on the graph with an asterisk *. The averages, standard deviations, and p values are presented in the SI.

All ions enhance the aggregation rate of aSyn, as evident when comparing the (Tris) Buffer only condition, “No Salt” to all other conditions. The Na^+^ and K^+^ ions both increase the aggregation rate, halving the t^lag^ and t^50^. However, the most distinct effect on aSyn aggregation is observed in the 2 mM CaCl^2^ condition, with a clear decrease in the aSyn nucleation time and an increase in the slope of the exponential elongation phase. The divalent Ca^2+^ ion speeds up the aggregation rate of aSyn to a much greater extent than the monovalent Na^+^ and K^+^ ions, even though its concentration is 75 times lower (2 mM vs 150 mM). In the presence of both Ca^2+^ and Na^+^ ions, the aggregation rate of aSyn is comparable to the rate in the presence of only Na^+^. These observations, in combination with the predicted absence of a structured binding site for Na^+^ and K^+^ ions ^58^, indicates that the different ionic conditions affect the aSyn kinetics differently. We have hypothesized that the binding of Ca^2+^ at the C-terminus modulates the release of long-range interactions of the aSyn monomer, while the Na^+^, and K^+^ ions have a non-specific, electrostatic effect on the monomer ^58^. To determine whether our observations from the kinetic assays can be attributed to structural changes in the aSyn monomer requires further studies. We thus decided to analyze the aSyn monomer structure and dynamics with a panel of biophysical techniques: ^1^H-^15^N HSQC NMR, the cutting-edge millisecond HDX-MS, and SANS. In particular, we focus our efforts on disentangling the distinct effects of Ca^2+^ and Na^+^ ions on aSyn.

### The conformational ensemble of monomeric aSyn is influenced to various degrees by the presence of the different ions

Previously ^52^, we have studied the binding of Ca^2+^ ions to aSyn via HSQC NMR. Here, we extend this study to probe the aSyn monomer structure in the presence of 4 mM NaCl, 150 mM NaCl, and 150 mM NaCl with 4.2 mM CaCl^2^ (**Figure 2 a-c**).

**Figure 2:**
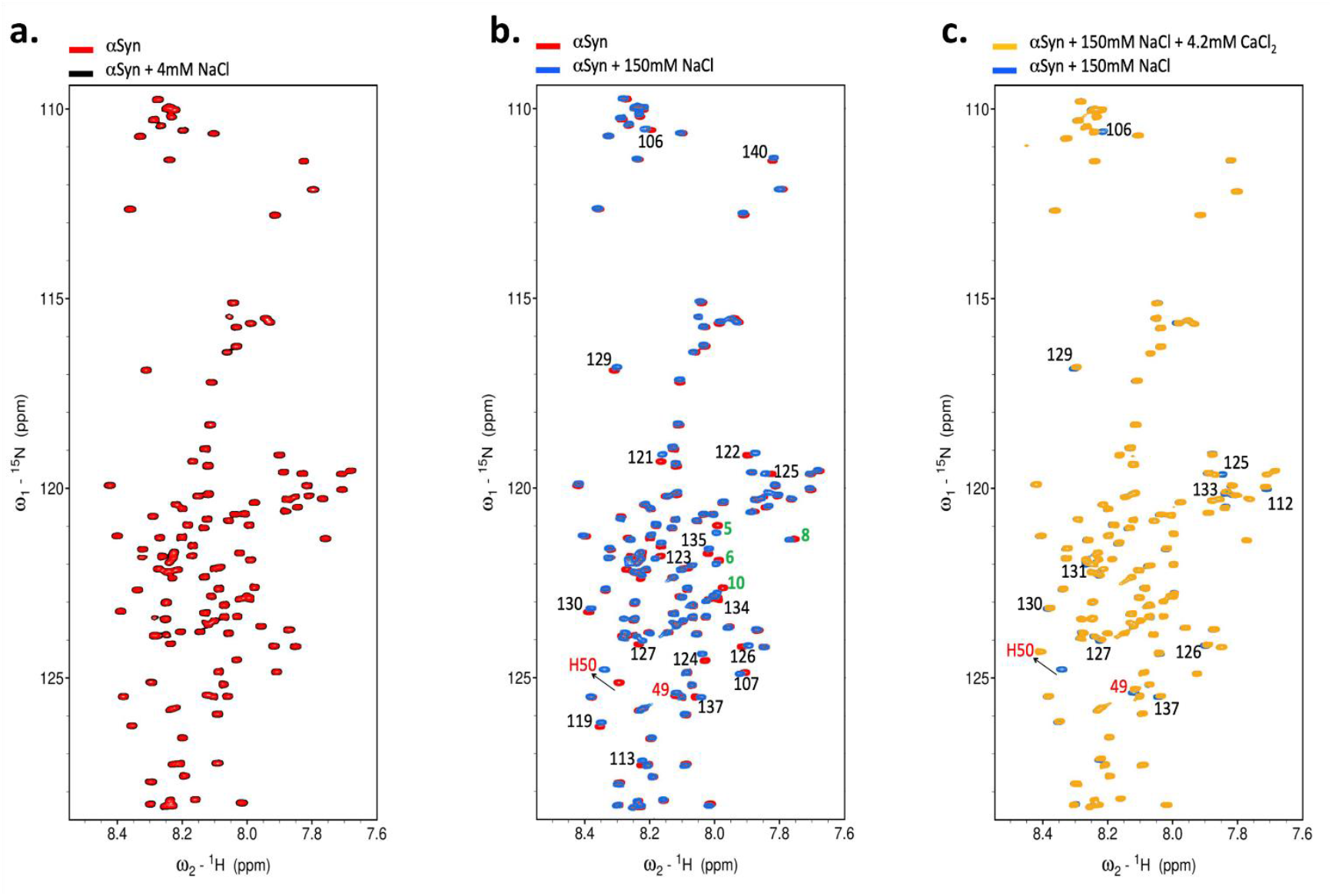
^1^H-^15^N HSQC NMR shows CSPs across the sequence of monomeric aSyn in the presence of NaCl and specific residue interactions at the C-terminus in the presence of Ca^2+^. **a**. We do not observe chemical shift perturbations (CSPs) in the ^1^H-^15^N resonances of aSyn amide backbone upon the addition of 4 mM NaCl, **b**. The addition of 150 mM NaCl causes CSPs primarily at the C-terminus (e.g. residues 121, 122, 123 etc., labelled black) and at the N-terminus (e.g. residues 5, 6, 8, 10 etc., labelled green). These peak motions are attributed to the weakening of the electrostatic interactions between the N- and C-terminal regions, as well as to possible rearrangements within the C-terminal region (where repulsion between negative charges diminishes due to the higher ionic strength). We also observed CSPs for H50 and the neighboring residue 49 (labelled red). These CSPs are attributed to alterations in the pKa (from 6.8 to 6.5) due to the change in salt concentrations (from 4 – 150 mM) ^59^, **c**. Comparison of aSyn in 150 mM NaCl and 150 mM NaCl + 4.2 mM CaCl_2_ shows CSPs at C-terminal residues (arrows with assigned amino acid residues) likely associated to direct/specific binding to Ca^2+^.

We have previously observed chemical shift perturbations (CSPs) at the C-terminus upon calcium binding (residues 104, 107, 112, 119, 123, 124, 126, 127, 129, 130, 135, 136, and 137), which correlate with the expected calcium-binding site. In contrast, 4 mM NaCl (**Figure 2a**) does not induce any CSPs on the aSyn monomer spectrum, showing that the identified Ca^2+^ effect at the C-terminus is specific to the ion and can be attributed to a specific binding site. At 150 mM NaCl (**Figure 2b**), CSPs can be observed at C-terminal residues (labelled in black, 106, 107, 113, 119, 124, 127, 129, 130, 134, 135, 137, 140) and at certain N-terminal residues (labelled in green, 5, 6, 8, 10). These peak motions can be attributed to a charge effect, a weakening of the intramolecular electrostatic interactions between N- and C-terminal regions, as well as to possible rearrangements within the C-terminal region, where repulsion between negative charges diminishes due to the high ionic strength. Intriguingly, we also observe peak changes at H50 and the neighboring residue 49 (labelled red), as a result of the alteration in pKa (from 6.8 to 6.5) due to the change in salt concentrations (from 4 – 150 mM) ^59^. In the presence of 150 mM NaCl and 4.2 mM CaCl^2^ (**Figure 2c**), the CSPs previously recorded at the C-terminus (106, 112, 125, 126, 127, 129, 130, 131, 133, 137) are still identifiable with respect to the 150 mM NaCl spectrum, proving that the binding site is retained in the presence of NaCl, and thus Ca^2+^ binding still occurs at the C-terminus even in the presence of 150 mM NaCl.

We next employ SANS to characterize conformational changes and radii of gyration of aSyn monomers in the different ionic conditions. Monomeric aSyn is measured in buffer only, in the presence of 2 mM CaCl^2^, 150 mM NaCl, and 150 mM NaCl mixed with 2 mM CaCl^2^, and the radius of gyration of the protein is calculated for each condition via an ensemble optimisation method (EOM) (**Figure 3**). In the absence of salts (“No salt condition”), aSyn samples display the widest distribution of R^g^. In the presence of Ca^2+^, aSyn samples show two main conformational spaces: a more compact structure ∼ 30 Å and a larger structure ∼ 60 Å. aSyn in NaCl exhibits a broader conformational space ∼40-50 Å while aSyn in the presence of both Na^+^ and Ca^2+^ has a larger R^g^ between 50-60 Å. The presence of Ca^2+^, in the absence and presence of Na^+^, shifts the conformation to favor more extended structures compared to the “no salt” conditions. These results suggest that these cations may facilitate desolvating aSyn in water, thus promoting aggregation of the protein.

**Figure 3:**
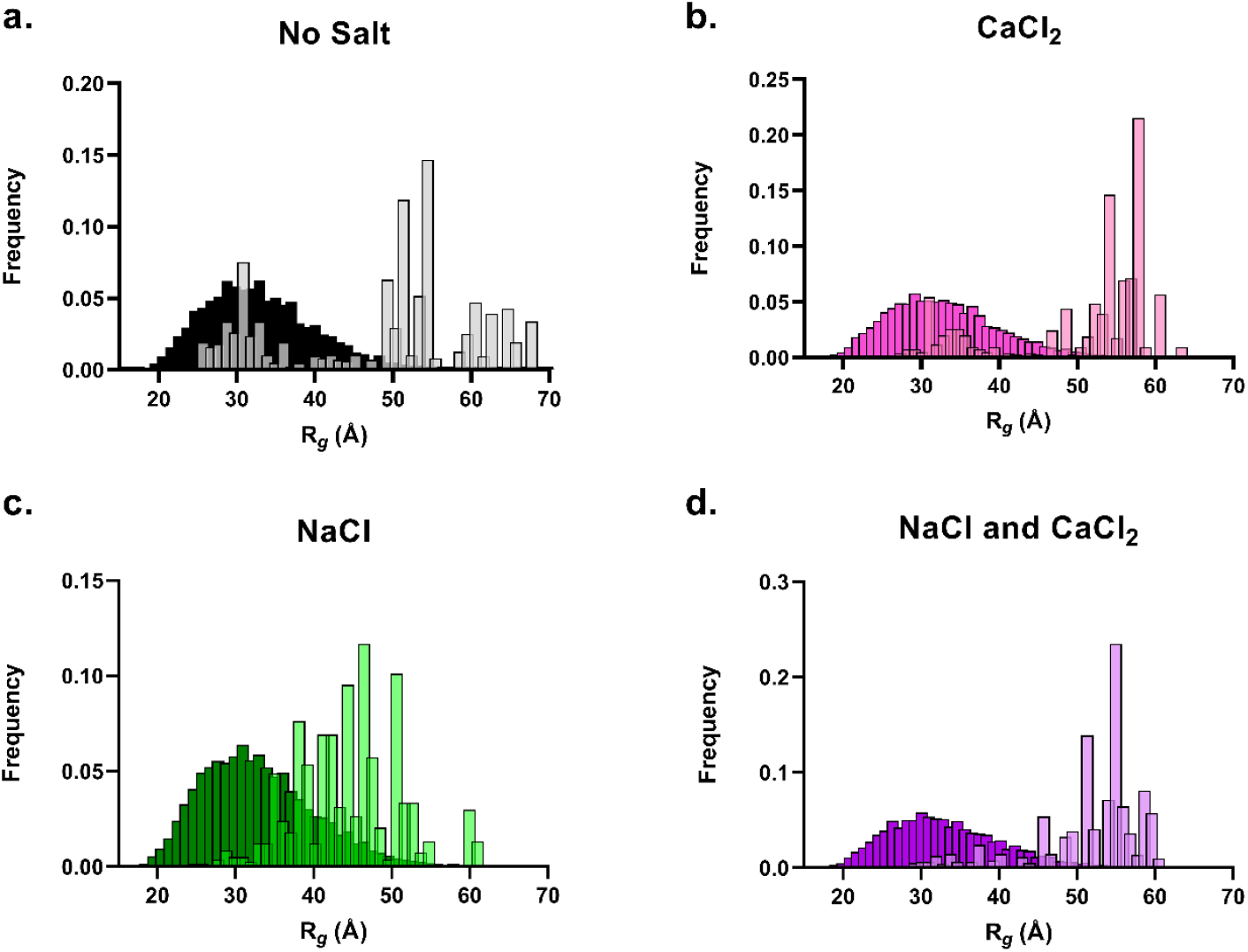
Small angle neutron scattering data show that Na^+^ induces an average extension of the conformation population of aSyn, whereas Ca^2+^ promotes a more extended conformation population. A pool of 10000 independent models (dark bars in each graph) based upon sequence, structural information (*i*.*e*., no defined structure for an IDP) is generated. The predicted scattering intensity from the models is compared to the experimental data and the 50 models of the best fit to the experimental data are selected as the most accurate representations (light bars in each graph). **a**. aSyn in the no salt tris buffer samples has the widest distribution of R_g_ (grey bars). **b**. aSyn in the CaCl_2_ buffer samples two main conformational spaces, a more compact structure ∼ 30 Å and a larger structure ∼ 60 Å (pink bars). **c**. aSyn in NaCl samples a more averaged size of conformational space ∼40-50 Å (green bars). **d**. aSyn in the presence of both NaCl and CaCl_2_ has larger Rg mostly between 50-60 Å (purple bars).

Having identified the differences between binding of Ca^2+^ and Na+ on aSyn via NMR, and the changes in the overall shape of the protein via SANS, we set out to capture the more localized and faster–changing protein dynamics of the aSyn monomer. We thus use millisecond HDX-MS with soft fragmentation via electron transfer dissociation (ETD) a technique that can be used to measure the dynamics of intrinsically disordered proteins at a very high temporal (subsecond mixing) and structural (ETD fragmentation) level.

The aSyn monomer is incubated in deuterated buffer (D_2_O), for timepoints ranging from 50 ms to 30 s in different conditions (Buffer only, 2 mM CaCl_2_, 150 mM NaCl, 150 mM NaCl and 2 mM CaCl_2_). It should be noted that the HDX rate is itself affected by the ionic strength of the buffer. For this reason, uncorrected deuterium uptake consists of the sum of the contributions of the ions to the HDX chemical exchange rate, and the changes in HDX due to differences in the conformation of the protein (which we set out to probe).

The data are thus corrected for the contribution of environmental conditions to the HDX chemical exchange rate, using the model disordered peptide, bradykinin, in the exact same buffer conditions as aSyn, as previously described ^53,60^. Therefore, we are able to make direct comparisons between the effect of the ions on the aSyn conformational ensemble. The HDX-MS data are plotted as the difference in deuterium uptake between protein states, for each labelling time point (50 ms to 30 s) across the protein sequence (**Figure 4**). This results in a deuterium uptake heatmap, in which positive values (blue) indicate that protein state 1 is deprotected compared to protein state 2, and negative values (red) indicate that state 1 is protected compared to state 2.

**Figure 4:**
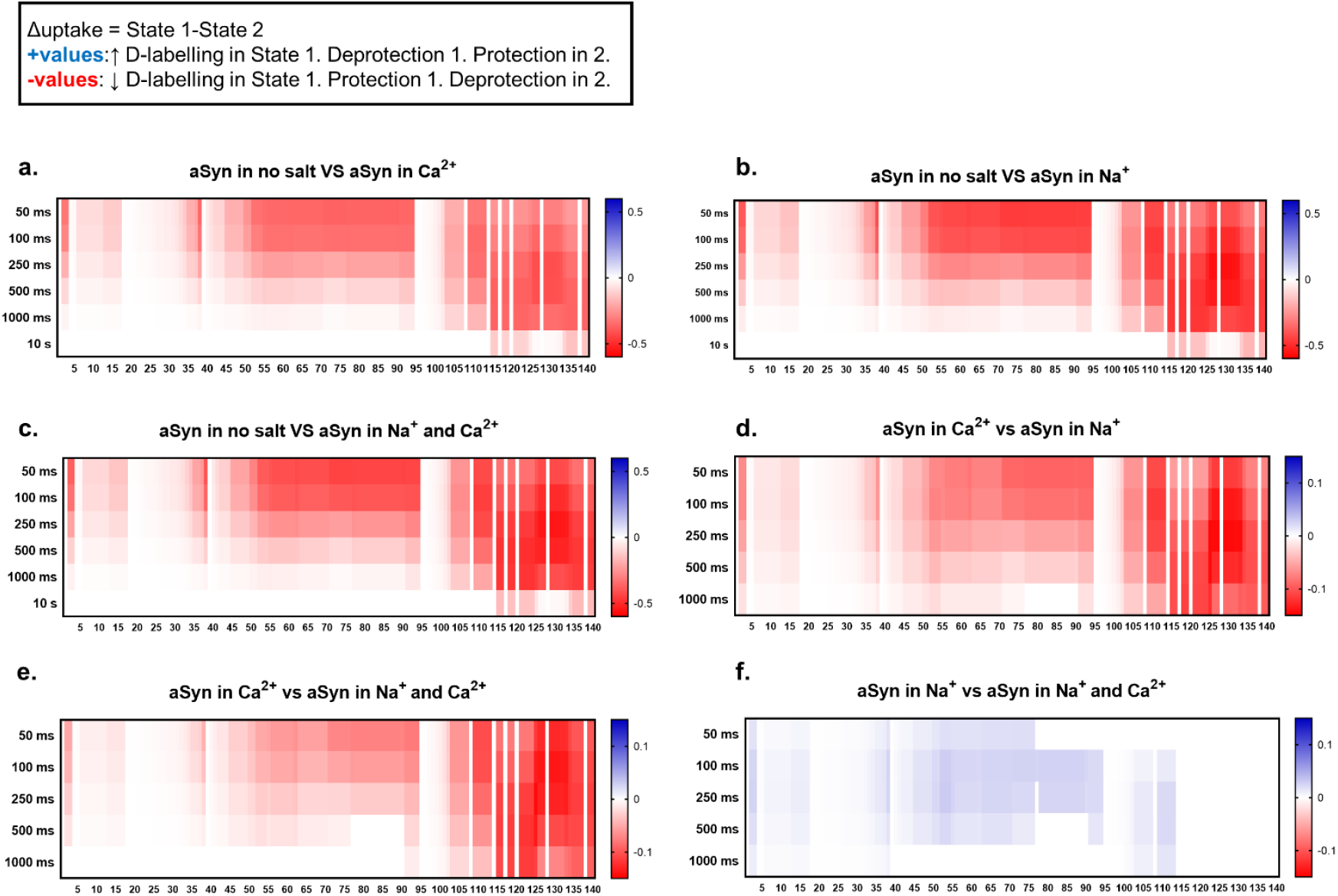
HDX-MS reveals that all ions induce a deprotection of monomeric aSyn, with Na^+^ having the strongest effect. Heatmaps showing significant differences (nonwhite) in deuterium uptake per timepoint during an on-exchange reaction between STATE 1–STATE 2 (*e*.*g*., aSyn in no salt vs aSyn with Ca^2+^). The conditions studied are aSyn in 20 mM Tris pH 7.4, with addition of 2 mM CaCl_2_, 150 mM NaCl, 150 mM NaCl and 2 mM CaCl_2_. X-axis: Protein sequence, y-axis: HDX labelling time point (50 ms – 10 s). Positive values are in red and represent decreased uptake in STATE 2, whereas negative values are in blue and represent increased uptake in STATE 2. Increased uptake indicates more solvent exposure and/or less participation in stable hydrogen-bonding networks. **a**. Monomeric aSyn is more protected when no salt is present compared to the addition of Ca^2+^, **b**. the addition of Na^+^, **c**. the addition of Na^+^ + Ca^2+^ **d**. Monomeric aSyn in Ca^2+^ is more protected compared to in Na^+^. **e**. aSyn in Ca^2+^ is more protected than in Na^+^ and Ca^2+^ and **f**. aSyn in Na^+^ is less protected than in Na^+^ and Ca^2+^. Data analysis was performed in DynamX (Waters) and hybrid significance testing was performed using Welch’s t-test (p-value of 0.05) and global significance thresholding.

When comparing the deuterium uptake of the “No salt buffer only” condition to all other conditions (aSyn + Ca^2+^, aSyn + Na^+^, aSyn + Ca^2+^ + Na^+^), aSyn is in its most protected/compacted conformation in the “No salt buffer only” condition, indicating that all tested ions cause deprotection (**Figure 4 a-c**). This deprotection is most pronounced at the NAC region residues 60-90 and residues 120-135 of the C-terminus. A binary comparison of aSyn deuterium uptake in the presence of Ca^2+^ versus in the presence of Na^+^ (**Figure 4 d**) shows that aSyn in the Na^+^ state is more deprotected/exposed, particularly at residues 60-90, 109-112, 120-125, and 130-135. A similar conclusion can be drawn when comparing aSyn in the Ca^2+^ condition versus the Na^+^ + Ca^2+^ condition (**Figure 4 e**). The (Na^+^ + Ca^2+^) state is more protected/compact than the Na^+^ state, particularly at residues 109-112,120-125, 130-135 of the C-terminus and residues 60-90 of the NAC region. To summarise, aSyn is in its most protected state when there are no salt ions present, is moderately deprotected in the presence of Ca^2+^, more deprotected when Na^+^ and Ca^2+^ is added, and most deprotected in Na^+^ only (Deuterium uptake: Na^+^> Na^+^ + Ca^2+^> Ca^2+^> No salt).

Overall, the information from the biophysical studies of the aSyn monomer structure in response to the different ions can be summarized in a model as follows: aSyn in buffer only (no salt, no Ca^2+^) inhabits a compact conformational ensemble. The addition of Na^+^ biases the conformational ensemble to more extended, deprotected structures (higher R_g_ at ∼40-50 Å, higher deuterium uptake), with no specific binding site (CSPs across the sequence at the N-terminus and C-terminus). The addition of Ca^2+^ leads to a more moderate conformational extension throughout the sequence (higher deuterium uptake, with two sub-populations at R_g_ at ∼ 30 Å and ∼ 60 Å), with localised perturbations at the C-terminus (CSPs at C-terminal residues), at the expected Ca^2+^ binding site. In the presence of both ions, the calcium binding site is retained (CSPs detected at the C-terminus) and the conformational ensemble is biased towards more extended structures which are less extended than the Na^+^ only condition, but more extended than the Ca^2+^ only condition.

To further validate our interpretation of the biophysical data, and to relate the monomer observations back to the aggregation rate of aSyn, we used Molecular Dynamics (MD) simulations to model the protein in the distinct local environments it encounters, to compare modelled aSyn monomeric to experimentally observed structured using ensemble techniques but, importantly, also to study the influence of the hydration shell of aSyn in the presence of the different ions.

### Modelling the aSyn conformation ensemble in distinct ion environments confirms structural changes observed by other biophysical techniques and reveals that Ca^2+^ significantly slows down water mobility in the hydration shell of aSyn

We used MD simulations to model aSyn monomer structures in conditions with minimum salt ions, 150 mM Na^+^, 34 Ca^2+^ ions, and a combination of 150 mM Na^+^ and 34 Ca^2+^ ions. In each case, the system was neutralized with Cl^-^ ions. We have chosen 34 Ca^2+^ ions as aSyn acts as a Ca^2+^ scavenger, and also because we want enough Ca^2+^ ions to saturate the binding site while allowing for approximately 10 mM CaCl_2_ left in solution. We extract monomers from PDB structures 2n0a (resolved by ssNMR), 8ads (resolved by cryoEM) and 8adw (resolved by cryoEM) as starting conformations in the simulations. Missing residues in 8a9l and 8ads are replaced by rigid body splicing of residues present in 2n0a using PyMOL. The simulations are run for over 1.5 µs at 303.15 K and we observe the conformational changes of the monomer throughout the simulation window as well as the ion contacts to the protein. Further to that, we observe the water molecules in the hydration shell of the protein in fine time-point simulations, where each solvent molecules’ total displacement is measured every 5 ps over a period of 3 ns, allowing the study of the solvent behaviour around the protein in the different environmental conditions.

By observing the Na^+^ and Ca^2+^ persistence times on the aSyn monomer (**Figure 5a,b**), we can infer that the binding of Ca^2+^ to aSyn is tighter, while Na^+^ ions have only transient interactions with the protein. This strengthens our previous experimental observations that indicate that the Na^+^ ions are simply adducts on the aSyn monomer. To disentangle the conformational dynamics of the protein in the different conditions, we plot the distances sampled between different areas of the protein in response to the added ions (**Figure 5c-e**). Overall, the addition of Na^+^ pushes the distribution towards slightly more elongated structures, particularly increasing the distance between the NAC region and the C-terminus. The Ca^2+^ ions also bias the conformational ensemble towards more elongated structures, increasing both the distance between the N-terminus and the NAC region as well as the distance between the NAC region and the C-terminus.

**Figure 5:**
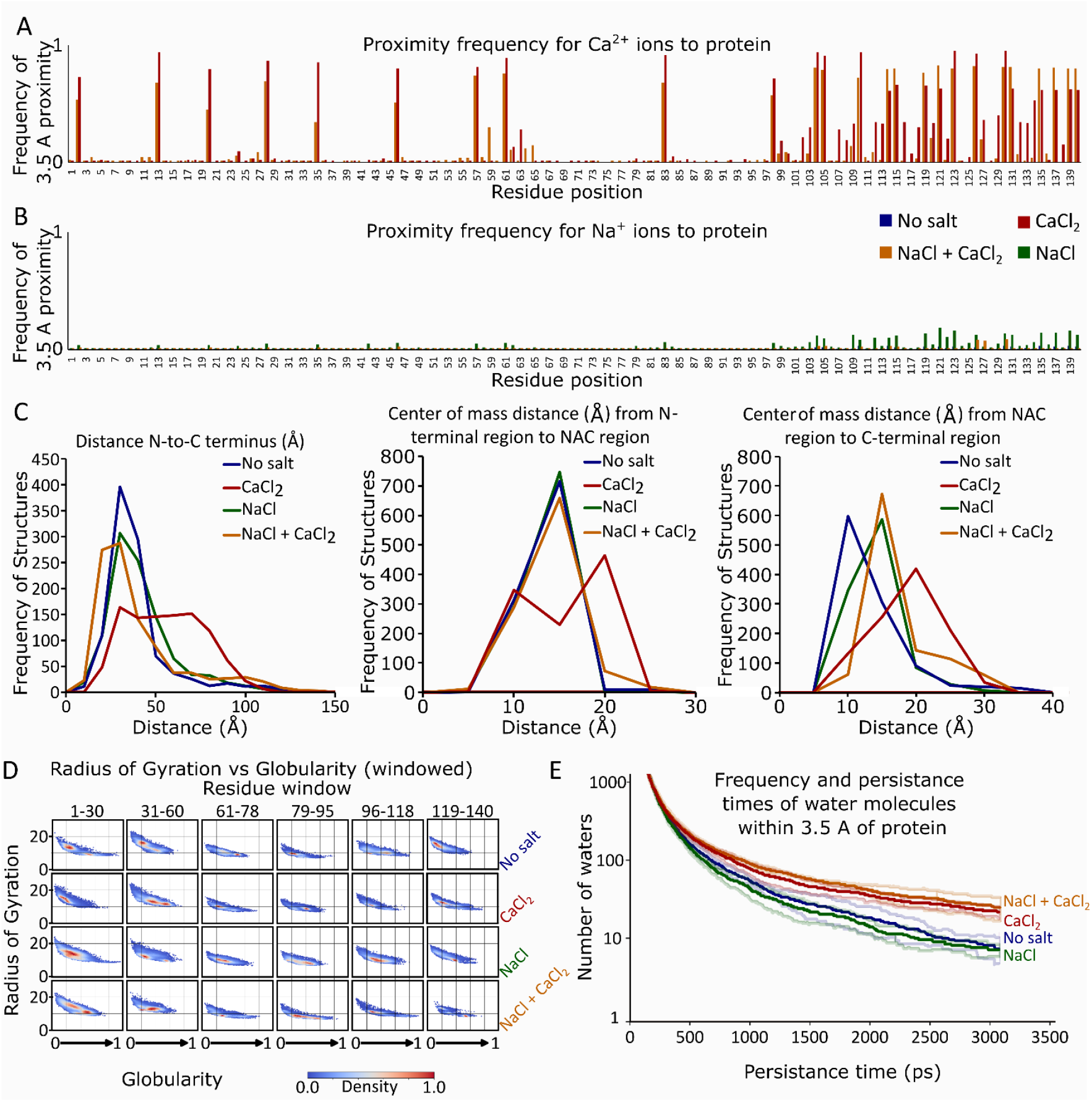
MD Simulations on monomeric aSyn in different ionic conditions confirm experimental results and reveal that Ca^2+^ ions have a higher aSyn persistence time compared to Na^+^ and that Ca^2+^ reduces solvent mobility around aSyn. **a.-b**.: Proximity frequency of Na^+^ and Ca^2+^ ions to the aSyn monomer. **c.-e**.: Distance of the N-terminus to the C-terminus, N-terminus to NAC region and NAC region to C-terminus in different ionic conditions, indicating conformational changes. **f**.: Radius of gyration (R_g_) plotted against protein globularity across the sequence (six residue windows) in different ionic conditions, **g**.: Persistence time of water around aSyn in different ionic conditions. The intention of the simulations was to restrict, satiate and saturate the calcium binding sites. As such, simulations were carried out under the following conditions; ‘No Salt’ contained the minimum ions required to equilibrate the system (10 Na^+^), ‘Unsaturated CaCl_2_’ (20 Ca^2+^ ions), ‘CaCl_2_’ (34 Ca^2+^ ions), ‘NaCl and CaCl_2_’ (150 mM Na^+^ and 34 Ca^2+^ ions), ‘NaCl’ (150 mM Na^+^) and ‘Saturated CaCl_2_’ (150 mM Ca^2+^), in each instance Cl^-^ ions were used to equilibrate the charge.

To gain further information on the shape of the protein and its compaction/extension, we calculate and plot the radius of gyration, *R*_*g*_, versus the globularity of the protein for each of its primary sequence regions (N-terminus, early 1-30 and late 31-60; NAC region, early 61-78 and late 79-95; and C-terminus early 96-118 and late 119-140) (**Figure 5d**). Overall, when comparing the aSyn monomer (full length) in the “No salt” condition to the ion conditions, we infer that the Na^+^ ions do not drastically alter the conformational ensemble, whereas the Ca^2+^ ions shift the population towards more extended (higher R_g_ values) and less globular (lower globularity values) structures, and a mix of the two ions slightly shifts the population towards more globular structures (**Figure 5d**). More specific information can be extracted for different areas of the protein: Na^+^ pushes towards more extended and less globular conformations at the early N-terminus (1-30), and towards more globular and less extended conformations at the NAC region and the C-terminus. The Ca^2+^ ions cause an extension and thus a decrease in globularity at the N-terminus (particularly at the early N-terminus, 1-30), have no significant effects at the NAC region, and cause a concurrent decrease in globularity at the early C-terminus as well as an increase in globularity at the late C-terminus, perhaps indicating a structural rearrangement near the binding site. A mix of the two ions push towards an extension of the N-terminus, and an increase in globularity at the NAC region and the C-terminus. Overall, we can see that both ions induce extension at the N-terminus, while most of the differences between Ca^2+^ and Na^+^ can be traced to the C-terminus, at the Ca^2+^ binding site. These observations correlate well with our experimental biophysical data (NMR, HDX-MS, SANS) on the aSyn monomer.

To further explain our observations in the bulk aggregation kinetics experiments, we next focus on the hydration of the protein in different conditions, by assessing the persistence time of water molecules around the aSyn monomer (**Figure 5e**). The highest values of water persistence are observed in the presence of Ca^2+^, while the addition of Na^+^ does not have such a dramatic effect on water mobility in the solvation shell. We observe that the presence of Ca^2+^ slows down the solvent in the hydration shell of the aSyn monomer, which correlates well with the increased aggregation propensity observed in the kinetics experiments.

### Distinct fibril polymorphs are formed in the presence of the different ions

To determine whether the detected differences in monomer conformation correlate with the formation of distinct fibril polymorphs, the structure of the fibrils at the end of the kinetic assays is probed via Atomic Force Microscopy (AFM). In order to quantitatively assess the morphological features of the various species, measurements of fibril height (h) and periodicity (p) are acquired (**Figure 6 a-d**).

**Figure 6:**
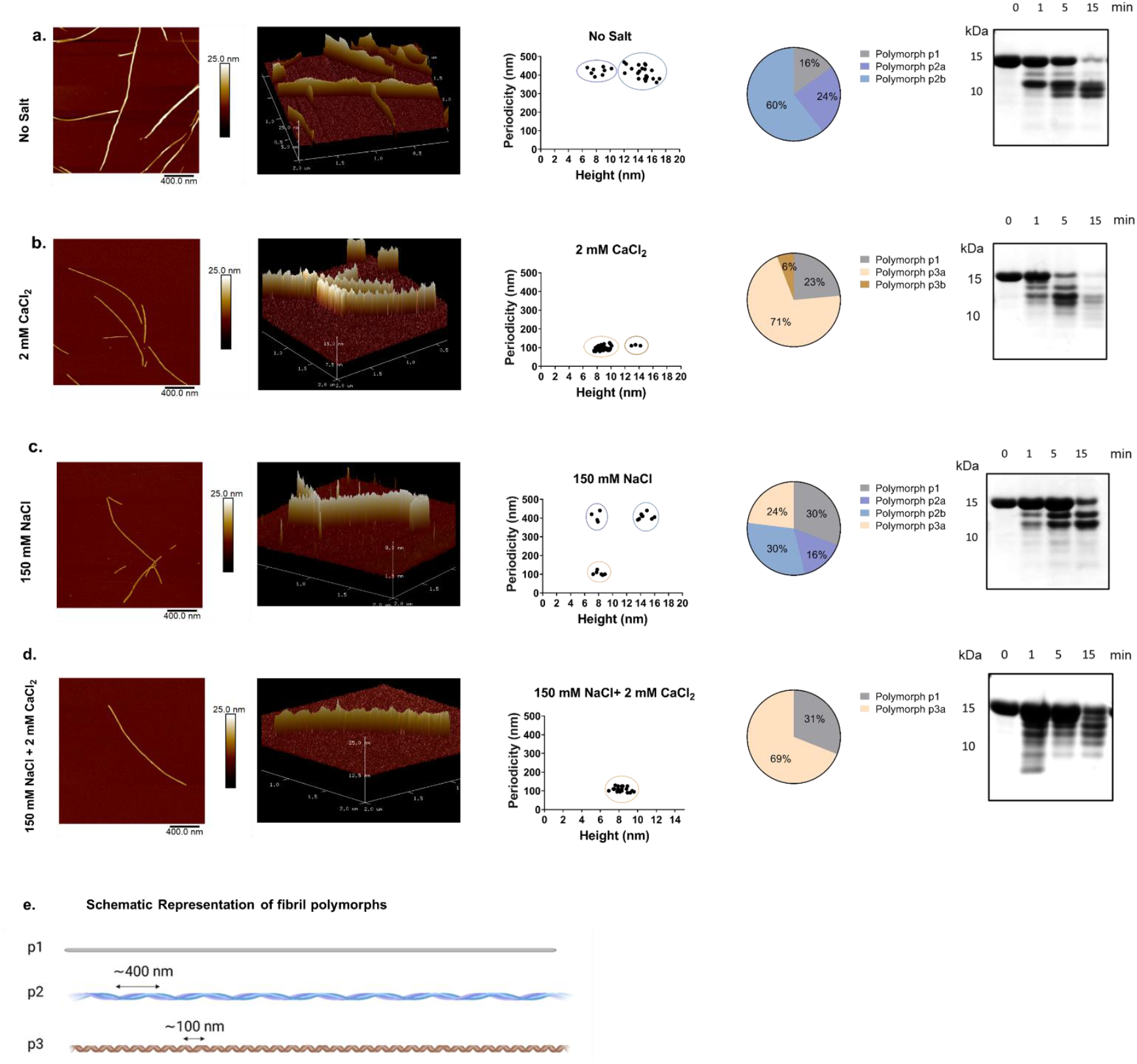
Different ionic conditions induce distinct aSyn polymorphs with Ca^2+^ inducing more twisted fibril structures as shown by AFM. **a.-d**.: Fibrils formed in no salt conditions, with 2 mM CaCl_2_, 150 mM NaCl, 150 mM NaCl and 2 mM CaCl_2_, respectively. First column: representative AFM image of the sample. Second column: 3D representation of the same image, Third column: Scatter plots of fibril periodicity and height. The heights of the fibrillar species were obtained by averaging measurements from individual fibrils (∼ 4 points per individual fibril). Fibril periodicity was determined by sectioning along the length of a fibril and averaging the distance measured between two adjacent peaks in height. The different polymorph populations are indicated with circles on the scatter plots. Polymorph p1 is not plotted on the scatter plot as it has no periodicity. Fourth column: Relative abundance (%) of each fibril polymorph in the sample. Fifth column: limited proteolysis of fibrils with proteinase K. Samples were incubated for four time points 0, 1, 5, 15 min and run on an SDS-PAGE gel. **e**.: Schematic representation of the three fibril polymorph populations identified across the samples. AFM images were collected in ScanAsyst Fluid tapping mode, with a resolution of 512 lines with a scan rate of 1.5 Hz. Fibril analysis was performed using NanoScope Analysis v1.9 software.

In all conditions, a percentage of the total fibril population is nonperiodic, contains rod-like fibrils, with h= 7.5 ± 1.7 nm, which we term polymorph p1. In the “No salt” condition, two more fibril polymorphs are detected: polymorph p2a, (h= 8.6 ± 1.4 nm and p= 422 ± 18 nm), with a height approximately corresponding to aSyn protofibrils and polymorph p2b (h= 13.8 ± 1.4 nm and p= 413 ± 33 nm), with a height corresponding to aSyn mature fibrils. Considering that the periodicity of the p2b fibril population (p2b) is approximately equal to the periodicity of the second population (p2a), it is very likely that the shorter fibrils of the p2a polymorph intertwine to form the mature tall fibrils of p2b.

The addition of Ca^2+^ biases the formation of more periodical, twisted fibrils. Polymorph p1 can still be seen in the sample, but the second, dominant population (relative abundance 75%) of fibrils is polymorph p3a (h=8.7 ± 0.7 nm and p=96 ± 15 nm). Some fibrils with the same periodicity (111 ± 5 nm) but increased height (13.5 ± 0.5 nm) are also detected and termed “p3b”. This again indicates that the fibrils p3a and p3b, which share the same periodicity, are the protofibrils and full fibrils of the same species. In the presence of Na^+^, aSyn forms fibrils that can be classified into four populations: the p2a and p2b polymorphs with the lower periodicity, previously seen in the “No salt” sample, some p3a fibrils with higher periodicity, as well as a number of non-periodical fibrils, p1. In the presence of Na^+^ and Ca^2+^, aSyn primarily forms polymorphs p3a, p3b (∼70% relative abundance), while some non-periodical fibrils, p1, can also be detected (∼30% relative abundance). The “Na^+^ + Ca^2+^” condition is therefore very similar to the “Ca^2+^” condition, as the main fibril population arising in both cases is the more twisted, periodical species of the polymorph p3a.

To further characterise the aSyn polymorphs, the fibrils are subjected to limited proteolysis with proteinase K (protK) and run on SDS-PAGE gels. This proteolytic treatment only affects fibril surfaces but not their amyloid core, which is preserved. Fibril polymorphs have been found to have strain-specific digestion patterns ^61^. Distinct digestion patterns can be observed for the studied ionic conditions (**Figure 6 a-d, right column**), indicating that the structural differences observed by AFM stem from distinct fibril core structures of aSyn.

## DISCUSSION

Studying the conformational dynamics of the alpha-synuclein monomer is fundamental for elucidating the molecular mechanisms underlying its aggregation, but also to pave the way for the development of effective therapies. By focusing on the monomer dynamics, we can identify transient structural motifs or regions with a propensity to adopt certain secondary structures which are ‘on path’ to oligomerization and aggregation. In the present study, we interrogate the effect of physiologically relevant ions, primarily Ca^2+^ and Na^+^, on the conformational ensemble of the aSyn monomer, and thus focus on the earliest stages of the aggregation process. We link our information on the monomer structure to its aggregation propensity and resulting fibril polymorphism. We have shown that the Ca^2+^ and Na^+^ ions impact the monomeric structure in distinct ways.

We have used a panel of biophysical techniques, each of which provide different nature of information, to probe the effect of the ions on aSyn. NMR generates high-resolution data on specific atomic interactions and local structural changes but cannot distinguish between extension or compaction of the monomer structure. SANS provides low-resolution, ensemble-averaged data about the overall shape and size of the protein in solution, offering insights into global conformation changes but might miss subtle local structural changes. HDX-MS provides information about the dynamics and flexibility of different regions of the protein and highlights regions with varying degrees of protection, indicating structural stability and dynamics, but has a lower spatial resolution compared to NMR. The orthogonality of these biophysical techniques, in addition to MD simulations, have allowed us to build towards a simple structural model of the effect of ions on the conformational ensemble.

To summarise, the Na^+^ ion shows a general effect of extension on the aSyn monomer structure (HDX-MS and SANS) and no structured binding site. MD simulations support these observations by demonstrating that Na^+^ facilitates an extended conformation, especially affecting the NAC region and the C-terminus. In contrast, Ca^2+^ has a specific binding site at the C-terminus, which is retained in the presence of Na^+^, and binding of Ca^2+^ to the monomer leads to a moderate extension of the N-terminus and NAC region (HDX-MS, SANS, MD). A schematic of the aSyn monomer in each condition can be found in **Figure 7**.

**Figure 7:**
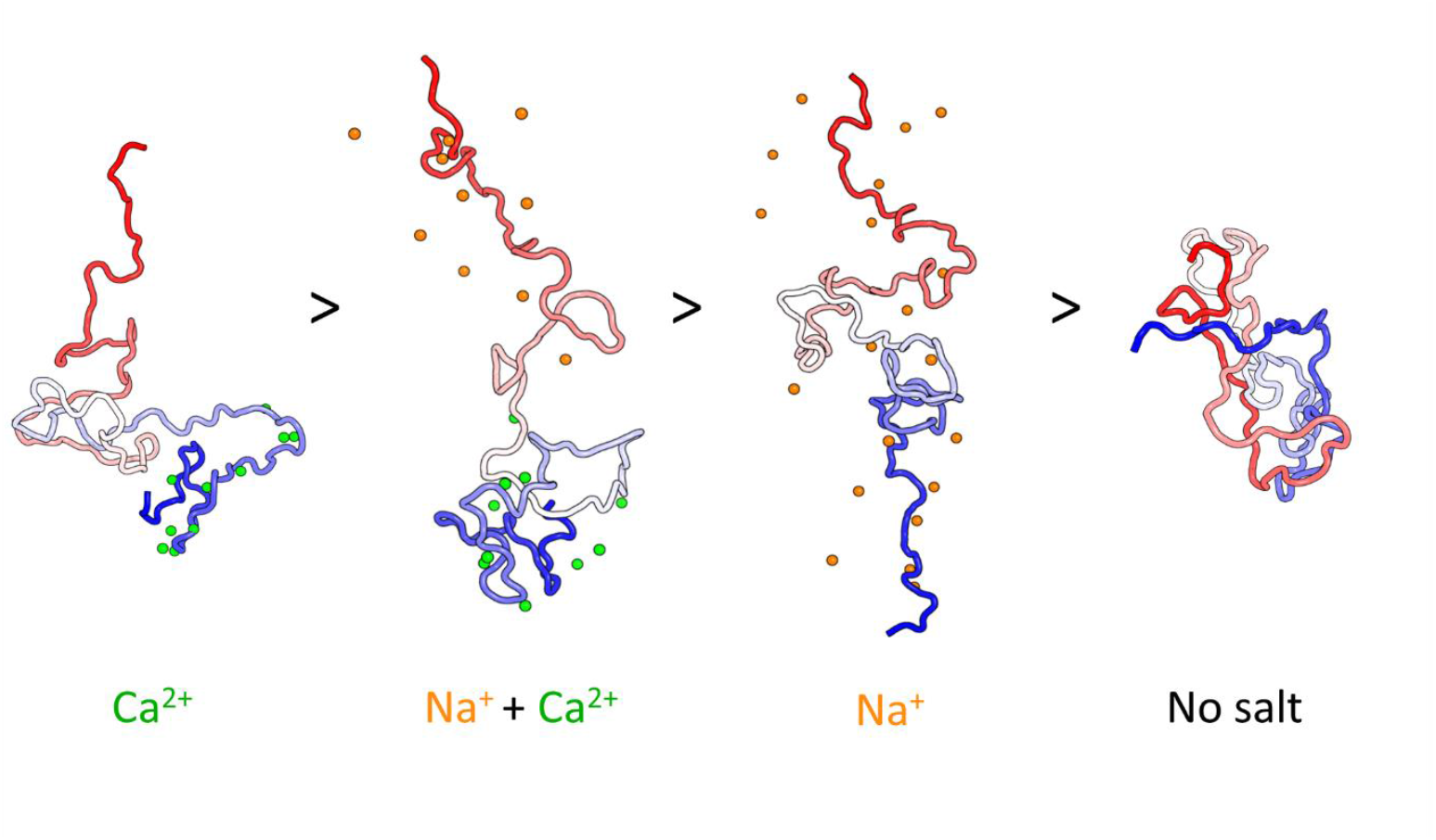
A proposed model for the aSyn conformational ensembles induced in distinct ionic environments, as informed by a combination of biophysical techniques and MD simulations, in reducing order of aggregation propensity. aSyn (N-terminus - red, NAC region - white, C-terminus – blue) in the presence of Na^+^ induces a general effect of extension on the aSyn monomer structure and does not contain a defined structured binding site on aSyn. In contrast, Ca^2+^ has a specific binding site at the C-terminus, which is retained in the presence of Na^+^, and binding of Ca^2+^ on the monomer structure and leads to a moderate extension of the N-terminus and the NAC region. In the absence of salt ions, the aSyn monomer samples a mostly compact conformation. Image created by manually adjusting a polyglycine structure. A 140AA polyglycine was generated using Alphafold2 and then manually edited with PyMol into different conformations.

It must be noted that we observe high persistence time of Ca^2+^ ions at the N-terminus (e.g. residues 2, 13, 20, 28) in the MD simulations, while in the NMR experiments the CSPs upon Ca^2+^ addition are localised at the C-terminus. We attribute this difference in information to the length of the simulations; aSyn is exposed to the ions for 1.5 μs, allowing the formation of ‘initial’ ensembles which may not represent the structural distributions in equilibrated ensembles. While the simulations may not have captured a ‘final conformational state’ they still highlight interesting ion and water effects even during these early conformations.

With regards to linking the conformational ensemble to the protein’s aggregation kinetics, we observe that an extension of the monomeric structure aids to increase the protein’s aggregation propensity (“no salt condition” compared to the ion conditions). This agrees with our previous observations ^52,53^, and multiple reports in the literature highlighting the effect of monomeric structure on aggregation kinetics ^52,62–66^. In our previous publication we have shown that the local ionic environment influences shifts in the aSyn conformational ensemble which are correlated with the aggregation kinetics, while here we have attempted a more sophisticated deconvolution of the individual ion contributions. The N-terminus is highlighted here as an area of particular importance. A recent study ^67^ has shown that deletion or substitution of the residues 36-42 at the N-terminus prevents aggregation, while three N-terminal truncations (deletions of 1-13-, 1-35, and 1-40 residues at the N terminus) have been shown to modulate both the aggregation kinetics, resulting in different fibril morphologies ^68^.

We propose that simply a bias towards more extended structures is not sufficient to explain the increase in aggregation propensity of the protein. That is because the most extended structures are observed in the presence of Na^+^, while the fastest aggregation kinetics are observed in the presence of Ca^2+^. We and others have shown that water plays an important role in protein aggregation ^54,69,70^, and decreased water mobility in the solvation shell correlates with an increased aSyn aggregation propensity ^54^. In this study, we observe increased water persistence times in the hydration shell in the presence of Ca^2+^ compared to Na^+^, via MD simulations. This correlates well with the aSyn aggregation kinetics in the presence of both ions (Na^+^ and Ca^2+^), which are intermediate compared the kinetics in the presence of the individual ions, as Na^+^ seems to balance the effect of Ca^2+^. We therefore suggest that the aggregation kinetics of monomeric aSyn is increased by a combination of conformational bias towards more extended structures, particularly at the N-terminus, together with a decreased water mobility in the solvation shell of the protein. Future experiments to further corroborate this claim will need to focus on investigating the mobility of the protein in the different ionic environments via ^1^H-^15^N HSQC NMR spectroscopy and Terahertz (THz) spectroscopy ^54^.

Regarding fibril polymorphism, AFM reveals distinct fibril morphologies formed under different ionic conditions. This finding suggests that the conformational biases induced by specific ions in the monomer state can lead to the formation of diverse fibril structures, which may correlate with the heterogeneous nature of Lewy bodies in PD. The conserved core of the amyloid fibrils has also been found to differ between polymorphs, as shown using the proteinase K degradation assay. The degradation pattern of the Na^+^ condition aligns well with a previously reported “finger-print” of a polymorph termed “ribbon” ^61^, while the digestion patterns of the polymorphs formed in the other conditions cannot not be identified in the literature. A difference in fibril polymorph toxicity has been reported in multiple studies ^71–73^ with fibrils formed *in vitro* in high salt concentrations forming more twisted periodical fibrils being more toxic. In a disease context, and on a molecular level, aSyn is likely to encounter higher Ca^2+^ concentrations in the extracellular space, especially due to neuronal damage or calcium dysregulation. Since aSyn can re-enter cells through endocytosis, it is conceivable that aSyn is consequently able to seed and template aggregation of endogenous aSyn, propagating the twisted fibril structures. A recent study on ex-vivo samples from post-mortem tissue has found variations in periodicity across different clinical phases of PD ^74^. The fibrils have displayed distinct periodic patterns influenced by the clinical stage, with shorter and more twisted fibrils arising in later PD stages, further suggesting a correlation between aSyn propagation, the ionic environment, fibril structure and disease progression.

Overall, our findings underscore the critical role of the ionic environment in regulating the structural dynamics and aggregation propensity of aSyn, as well as fibril polymorphism. By revealing how specific ions influence aSyn behavior, this study contributes to a deeper understanding of the molecular underpinnings of PD and other synucleinopathies and opens new avenues for therapeutic intervention by targeting aSyn, in particular in the extracellular space, before it can re-enter the neighboring neuron and enhance propagation.

## METHODS

### Thioflavin-T (ThT) Binding Assay in 96-Well Plates

Thioflavin – T (ThT) kinetic assays are used to monitor the aggregation of aSyn in different buffer conditions. For sample preparation, 20 μM (final concentration) of freshly made ThT solution (Abcam, Cambridge, UK) in distilled water is added to 50 μL of 20 μM, aSyn in 20 mM Tris pH 7.4, supplemented with 2 mM CaCl_2_ (“Ca”), 150 mM NaCl (“Na”), 150 mM NaCl and 2 mM CaCl2 (“Na + Ca”), or 150 mM KCl (“K”).

All samples are loaded in nonbinding, clear 96-well plates (Greiner Bio-One GmbH, Germany) which are then sealed with a SILVERseal aluminium microplate sealer (Grenier Bio-One GmbH). Fluorescence measurements are taken using a FLUOstar Omega microplate reader (BMG LABTECH GmbH, Ortenbery, Germany). Excitation is set at 440 nm and the ThT fluorescence intensity is measured at 480 nm emission with a 1300 gain setting. The plates are incubated with double orbital shaking for 300 s before the readings (every 60 min) at 300 rpm. Three repeats are performed with 6 replicates per condition. Each repeat is performed with a different purification batch of aSyn (biological replicate). Data are normalised to the well with the maximum fluorescence intensity for each plate and the empirical aggregation parameters t_lag_, t_50_, k, are calculated for each condition, based on the equation:

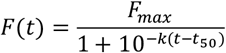

where *F* is the normalised fluorescence to the highest value recorded in the plate repeat, *F*_*max*_is the maximum fluorescence at the plateau, *k* is the slope of the exponential phase of the curve, and *t*_50_ is the time when 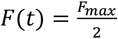.

One-way ANOVA is used to calculate statistical significance between samples using GraphPad Prism 8 (GraphPad Software, USA).

### SEC-HPLC (Size exclusion–High-performance liquid chromatography)

At the end of the ThT-based aggregation assays, the amount of remaining monomer of aSyn in each well is determined by analytical size exclusion chromatography with HPLC (SEC-HPLC). SEC analysis is performed on the Agilent 1260 Infinity HPLC system (Agilent Technologies, UK) equipped with an autosampler and a diode-array detector using a AdvanceBio SEC column (Agilent Technologies, UK) in 20 mM Tris pH 7.4 at 0.8 mL/min flow-rate. 25 μL of each sample is injected onto the column and the elution profile is monitored by UV absorption at 220 and 280 nm. The area under the peaks in the chromatogram of absorption at 280 nm is calculated to provide the monomer concentration. Samples for a standard curve spanning from 5 uM aSyn to 40 uM aSyn are run on the column, to relate the area under the curve to protein concentration.

### 1H-^15^N HSQC NMR

NMR experiments are carried out at 10 °C on a Bruker spectrometer operating at ^1^H frequency of 700 MHz equipped with triple resonance HCN cryo-probe and data are collected using the Topspin 4.4.0 software (Bruker, AXS GmBH). The ^1^H-^15^N HSQC experiments are recorded using a data matrix consisting of 2048 (t2, 1H) × 220 (t1, 15N) complex points. Assignments of the resonances in ^1^H-^15^N-HSQC spectra of aSyn are derived from our previous studies and data are analysed using the Sparky 3.1 software. The perturbation of the ^1^H-^15^N HSQC resonances are analysed using a weighting function in Eq 1:

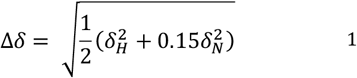

### Small-Angle Neutron Scattering (SANS)

Monomeric aSyn is isolated by gel filtration using a SuperdexTM 75 10/300 GL column (Cytivia, USA). The aSyn samples are buffer exchanged into D_2_O with 20 mM Tris pD 7.2 with the addition of either 150 mM NaCl, 2 mM CaCl_2_, or 150 mM NaCl + 2 mM CaCl_2_ using a HiTrapTM Desalting Sephadex G-25 column (#29048684, Cytivia). 200 uL of the sample solution is added to a 1 mm thick quartz cuvette (#QS, 100-1-40, Hellma, UK). Small angle neutron scattering (SANS) experiments [https://dx.doi.org/10.5291/ILL-DATA.8-03-974] are carried out on D22 at the Institut Laue-Langevin *-* The European Neutron Source, Grenoble, France. Samples are analysed at three sample–detector distances (2.5 m, 5.6 m and 17.6 m), with respective collimation lengths of 2.8m and 5.6m. The wavelength is 6 Å +/-10%. The source aperture is 40 mm x 55 mm, and the beam aperture is 7 mm x 10 mm. Samples are maintained at 10°C during the measurement. The raw counts from the detector for an empty cell and dark pattern are subtracted from the datasets. These are then scaled by the transmission and sample thickness as well as the direct flux measurement to obtain an absolute intensity. The data was reduced using Grasp^75^. Data recorded in different configurations are buffer-subtracted and stitched together into individual 1D plots of I(*q*) versus *q* for each sample using specialized macros in Igor Pro^76^.

### Ensemble Optimization Method (EOM)

The experimental SANS data are analyzed using ensemble optimization method (EOM) ^77^ (https://www.embl-hamburg.de/biosaxs/eom.html). The WT aSyn sequence and the SANS experimental data are used without a defined structure, due to aSyn being intrinsically disordered, to predict 10000 models in a RANdom Chains (RANCH) program. Genetic Algorithm Judging Optimisation of Ensembles (GAJOE) is then used to select from 10000 models, an ensemble of 50 models whose theoretical scattering intensity is most similar to the experimental scattering data. The pool of 10000 models and the 50 most predictive models are presented as size distributions in the form of the radius of gyration (R_g_ (Å)).

### Hydrogen-deuterium exchange mass spectrometry (HDX-MS)

For labelling times ranging between 50 ms and 5 min, hydrogen-deuterium exchange (HDX) is performed using a fully automated, millisecond HDX labelling and online quench-flow instrument, ms2min ^55,56^ (Applied Photophysics, UK), connected to an HDX manager (Waters, USA). For each cellular condition and three biological replicates, aSyn samples in the equilibrium buffer are delivered into the labelling mixer and diluted 20-fold with labelling buffer at 20 °C, initiating HDX. The duration of the HDX labelling depends on the mixing loops of varying length in the sample chamber of the ms2min and the velocity of the carrier buffer, calibrated to a precision of 1 ms. The protein is labelled for a range of times from 50 ms to 5 min. Immediately post-labelling, the labelled sample is mixed with quench buffer in a 1:1 ratio in the quench mixer to arrest HDX. The sample is then centered on the HPLC injection loop of the ms2min and sent to the HDX manager. For longer timepoints above 5 min, a CTC PAL sample handling robot (LEAP Technologies, USA) is used. Protein samples are digested onto an Enzymate immobilized pepsin column (Waters, USA) to form peptides. The peptides are trapped on a VanGuard 2.1 × 5 mm ACQUITY BEH C18 column (Waters, USA) for 3 minutes at 125 µL/min and separated on a 1 × 100mm ACQUITY BEH 1.7 μm C18 column (Waters, USA) with a 7-minute linear gradient of acetonitrile (5-40%) supplemented with 0.1% formic acid. Peptide samples do not require the initial peptic digestion step. The eluted peptides are analyzed on a Synapt G2-Si mass spectrometer (Waters, USA). An MSonly method with a low collisional activation energy is used for peptide-only HDX and an MS/MS ETD fragmentation method is used for HDX-MS-ETD. Deuterium incorporation into the peptides and ETD fragments is measured in DynamX 3.0 (Waters, USA).

### ETD fragmentation of aSyn peptides

The ETD reagent used is 4-nitrotoluene. The intensity of the ETD reagent per second, determined by the glow discharge settings, is tuned to give a signal of approximately 1e7 counts per second (make-up gas flow: 35 mL/min, discharge current 65 µA) to give efficient ETD fragmentation. Instrument settings are as follows: sampling cone 30 V, trap cell pressure 5e-2 mbar, trap wave height 0.25 V, trap wave velocity 300 m/s, transfer collision energy 8 V and transfer cell pressure 8e-3 mbar. Hydrogen-deuterium scrambling is measured using Peptide P1 under the same instrument conditions.

### HDX-MS Data analysis

The raw data are processed, and assignments of isotopic distributions are reviewed in DynamX 3.0 (Waters, USA). The post-processing analysis is performed using HDfleX ^60^. Briefly, the back-exchange-corrected data points for each peptide and ETD fragment are fitted using equation 2 in one-phase.

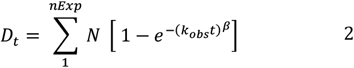

As the rate of HDX is affected by pH and ionic strength, which are not controlled in this study, it is crucial to normalize the solution effects between the different conditions being compared. Here, we use an empirical approach to normalization using the unstructured peptide bradykinin (RPPGFSPFR)^78,79^ to deconvolute the solution effects of the HDX from the protein structural changes. Due to the unstructured nature of bradykinin, all the differences in deuterium uptake seen from the different buffers can be assumed to be strictly from the changes in the chemical exchange rate effects, rather than from structural effects. The ETD fragments are combined with the peptide data using HDfleX ^60^ to give the absolute uptake information across the entire protein.

### Statistical significance analysis

The hybrid significance testing method along with data flattening is used here and is described elsewhere ^60^.

### Molecular Dynamics Simulations

Three aSyn models are generated from the PDB structures 2N0A, 8A9L and 8ADS. Missing residues in 8A9L and 8ADS are supplemented by residues from 2N0A. For each initial structure, six simulations are run with differing ionic constituents. Preliminary simulations (not described) show approximately 25 calcium binding sites per monomer. The intention of these simulations was to restrict, satiate and saturate the calcium binding sites, as such, simulations are carried out in the following conditions; ‘No Salt’ contained the minimum ions required to equilibrate the system (10 Na^+^), ‘Unsaturated CaCl^2^’ (20 Ca^2+^ ions), ‘CaCl^2^’ (34 Ca^2+^ ions), ‘NaCl and CaCl^2^’ (150 mM Na^+^ and 34 Ca^2+^ ions), ‘NaCl’ (150 mM Na^+^) and ‘Saturated CaCl^2^’ (150 mM Ca^2+^), in each instance Cl^-^ ions were used to equilibrate the charge. Input files are prepared with Amber Tools ^80^ packages tleap and parmed. Hydrogen atoms are added appropriate for pH 7. A truncated octahedron is solvated with Optimal Point Charges - OPC water[doi.org/10.1021/jz501780a] and ions appropriate for the relative investigation with a 14 Å gap from the protein to the periodic boundary. The system is parameterized using amber leaprc.protein.ff19SB, leaprc.water.OPC and the frcmod.ion-slm_1264_opc[https://doi.org/10.1021/acs.jctc.0c00194]f orcefields hydrogen mass repartitioning is used to distribute heavy atom mass onto neighboring hydrogen atoms, allowing 4 fs time-steps. Simulations are run using Amber22 on the Arc3 and Arc4 HPC at the University of Leeds, UK. Backbone protein atoms are restrained during 5000 steps of minimization and 500 ps of equilibration. Production runs are conducted for an average of 1.9 µs (20NA), 1.6 µs (8A9L) and 2.7 µs (8ADS) at 303.15 K, structures are written every 500 ps for analysis. A second set of fine time-scale simulations are performed using restart files from the above simulations at low RMSD areas of the trajectories for 3 ns at 303.15 K and structures are written every 5 ps for analysis. The ion proximity frequency, water persistence times and the distances between N and C-terminal and section center of masses are all calculated in PyMOL^81^. The radius of gyration is calculated in cpptraj (Amber Tools) using the ‘radgyr’ function. The globularity is calculated by taking the ratio of largest diagonalized eigenvalue and the smallest (Globularity = λ_x_/λ_z_), as calculated by the cpptraj function ‘principle’. Density maps and graphs are prepared in python using numpy, seaborn, matplotlib and scipy.

### Atomic Force Microscopy (AFM) analysis of fibril morphology

Fibrils formed at the end of ThT assays are analyzed by AFM. A freshly cleaved mica surface is coated with 0.1% poly-l-lysine, washed with distilled H_2_O thrice and dried under a stream of nitrogen gas. Samples from the microplate wells are then incubated for 30 min on the mica surface. The sample is washed thrice in the buffer of choice (for example, in 20 mM Tris, pH 7.4 for the Tris condition) to remove lose fibrils. Images are acquired in fluid using tapping mode on a BioScope Resolve AFM (Bruker, USA) using ScanAsyst-Fluid+ probes. 512 lines were acquired at a scan rate of 1.5 Hz per image with a field of view of 2-5 µm and for at least ten fields of view. Images are adjusted for contrast and exported from the NanoScope Analysis 8.2 software (Bruker). Measurements of fibril height and periodicity are performed by cross-sectioning across the fibril and across the fibril axis in NanoScope Analysis 8.2 software (Bruker). Statistical analysis of the height and periodicity measurements is performed in GraphPad Prism 8 (GraphPad Software, USA).

### Proteinase-K (ProtK) Limited Proteolysis of aSyn Fibrils

Fibrils formed in different conditions: aSyn in 20 mM Tris pH 7.4 (“No salt”), supplemented with 2 mM CaCl_2_ (“Ca”), 150 mM NaCl (“Na”), 150 mM NaCl and 2 mM CaCl2 (“Na + Ca”) were incubated at 37° C with Proteinase K (3 μg/ml) (Roche). Aliquots are removed at different time intervals following addition of the protease (0, 1, 5, 15 min) and transferred into Eppendorf tubes with proteinase K inhibitor phenylmethylsulfonyl fluoride (PMSF). Samples are placed at -80 °C for 3 h and dried using a freeze dryer and further solubilized by addition of pure HFIP(Hexafluoroisopropanol). After evaporation of HFIP, the samples are resuspended in gel loading buffer, heated for 10 min at 90 °C, and run on an SDS-PAGE (15%) gel.

## ASSOCIATED CONTENT

### Data Availability statement

The authors declare that the data supporting the findings of this study are available in this paper and its supplementary information files. Source data are provided with this paper. All mass spectrometry .raw files will be made available on request to JJP.

Code Availability statement: This study uses in-house developed software available to download: [http://hdl.handle.net/10871/127982] ^60^. Supporting information shows additional code used to calculate the Pearson correlation coefficients and to plot the correlation plots at each amino acid.

## Present Addresses

Maria Zacharopoulou: Department of Pharmacology, University of Cambridge, Cambridge, United Kingdom, Neeleema Seetaloo: Vertex Pharmaceuticals, Oxford, United Kingdom, Amberley Stephens: AstraZeneca, Cambridge, United Kingdom, Giuliana Fusco: Department of Pharmacy, University of Naples Federico II, Naples, Italy, Alfonso De Simone: Department of Pharmacy, University of Naples Federico II, Naples, Italy, Ioanna Mela: Department of Pharmacology, University of Cambridge, Cambridge, United Kingdom, Thomas McCoy: Delft University of Technology, Delft, Netherlands.

All authors have given approval to the final version of the manuscript.

## ABBREVIATIONS

aSyn: (Alpha-synuclein)
PD: (Parkinson’s Disease)
NMR: (Nuclear Magnetic Resonance)
HDX-MS: (Hydrogen-Deuterium Exchange Mass Spectrometry)
SANS: (Small-Angle Neutron Scattering)
MD: (Molecular Dynamics)
IDP: (Intrinsically Disordered Protein)
Rg: (Radius of Gyration)
SEC: (Size-Exclusion Chromatography)
ESI: (Electrospray Ionization)
TOF: (Time-of-Flight)
RMSD: (Root Mean Square Deviation)
DLS: (Dynamic Light Scattering)
HSQC: (Heteronuclear Single Quantum Coherence)
Chemical shift perturbations: (CSP)
PDB: (Protein Data Bank).

