## Supplementary Information for "Local ionic conditions modulate the aggregation propensity and influence the structural polymorphism of alpha-synuclein"

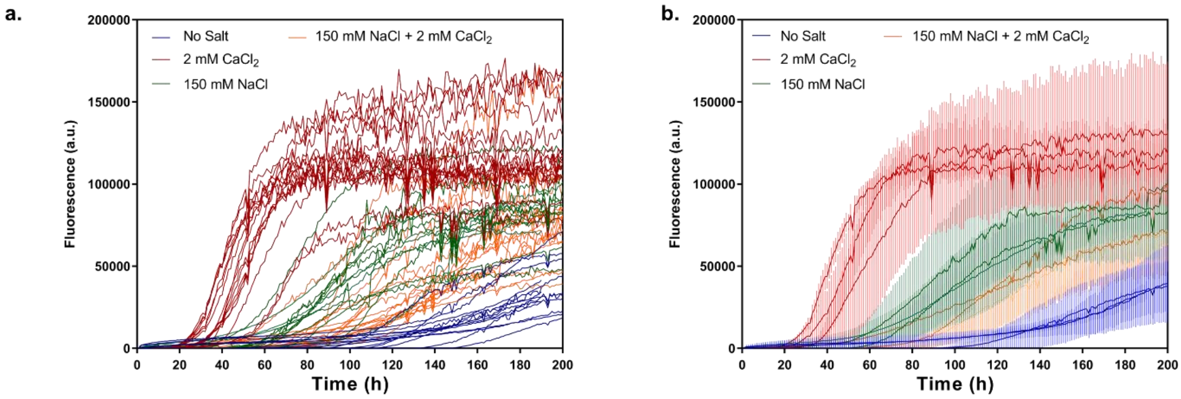


**Figure S1: (a) All individual ThT aggregation kinetics replicates across biological repeats and (b) average of traces for each biological replicate (N=3).** Fluorescence intensity plotted in arbitrary units (a.u.).

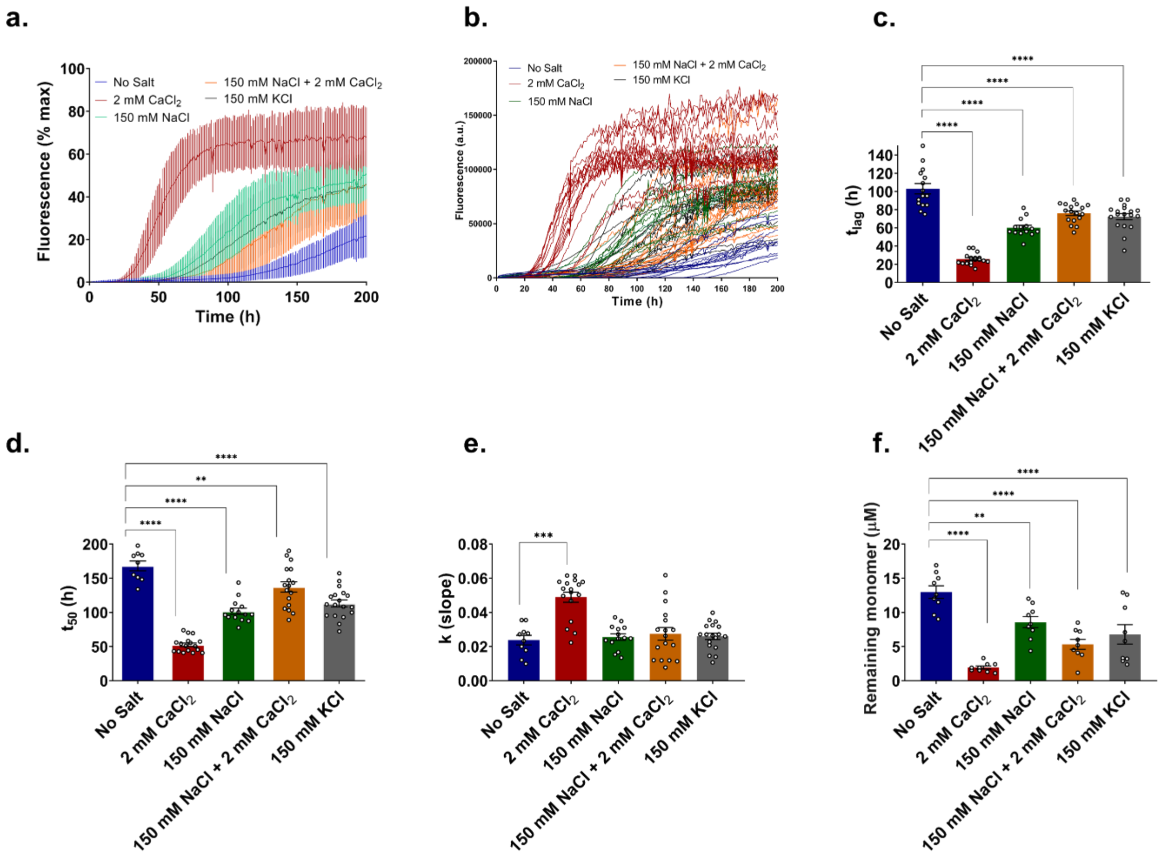


**Figure S2: ThT aggregation kinetics of WT aSyn is increased in the presence of all ions ($Na^+$, $Ca^{2+}$, $K^+$).** The conditions plotted here are aSyn in 20 mM Tris pH 7.4, with addition of 2 mM $CaCl_2$, 150 mM NaCl, 150 mM NaCl and 2 mM $CaCl_2$, 150 mM KCl. At least 6 replicates across three biological repeats were collected per condition. **a.** Kinetic traces. The average between traces of the same condition is shown in the graph and errors indicate 1 s.d., **b.** Each individual kinetic trace. Fluorescence intensity plotted in arbitrary units (a.u.) **c**. lag time ($t_{lag}$), **d**. time to reach 50 % of maximum aggregation ($t_{50}$) and **e.** the slope of the curve *k* were calculated by fitting Equation 1, and the mean plus error (1 s.d.) are displayed in the graphs. **f.** Remaining monomer concentration (µM) at the end of the aggregation assay was determined using SEC-HPLC. An ordinary ANOVA was used to calculate statistical significance between samples and significant differences are reported on the graph with an asterisk *.

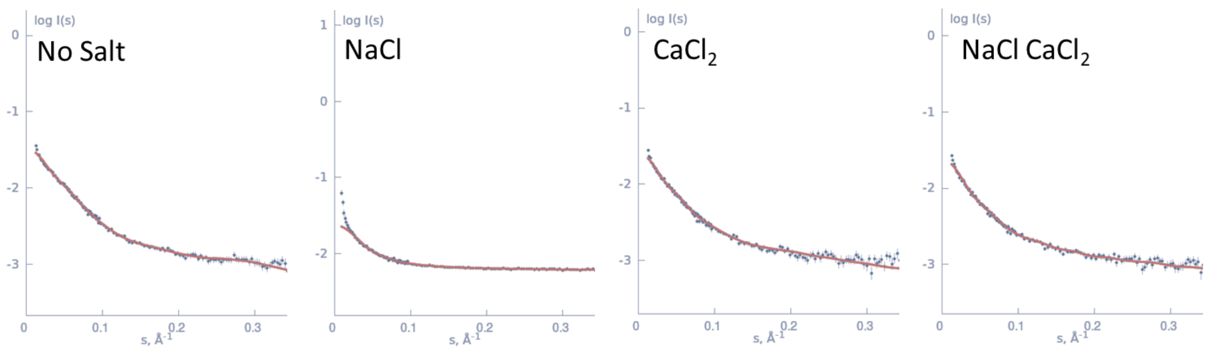


**Figure S3: SANS of aSyn in different ionic conditions.** GAJOE was used to fit to the best ensemble prediction from the data (red fitted line)

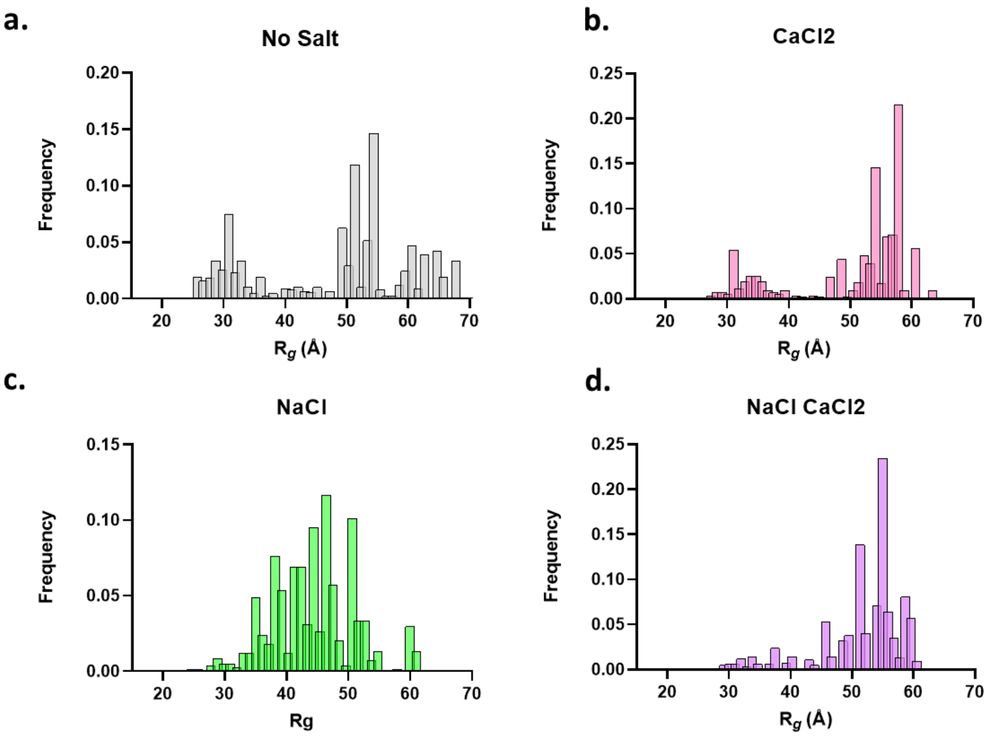


**Figure S4: Ensemble optimisation method (EOM) of SANS data shows varying distribution of the radius of gyration of aSyn in different ionic conditions.** A pool of 1000 independent models based upon sequence and structural information (i.e., no defined structure for an IDP) is generated.

The predicted scattering intensity from the models is compared to the experimental data and the 50 models of the best fit to the experimental data are selected as the most accurate representations, represented in the graphs.

Table S1: HDX-MS experimental technical details.

| **Data Set** | **aSyn** |
|---|---|
| HDX reaction details | 20˚C, pH 7.4<br>20 mM Tris<br>20 mM Tris + 2 mM $CaCl_2$<br>20 mM Tris + 150 mM NaCl<br>20 mM Tris + 150 mM NaCl + 2 mM $CaCl_2$ |
| HDX time course (ms) | 50, 100, 250, 500, 1000, 10000, 30000, 300000. |
| Back-exchange (mean / IQR) | 39.3% / 15.1% |
| Number of Peptides | 30 |
| Sequence coverage | 100% |
| Peptide Redundancy | 3.87 |
| Replicates (biological or technical) | 1 (biological), 3 (technical) |
| Significant differences in HDX (delta HDX > X Da) | 0.35 (95% CI, quartiles method of outlier removal) |

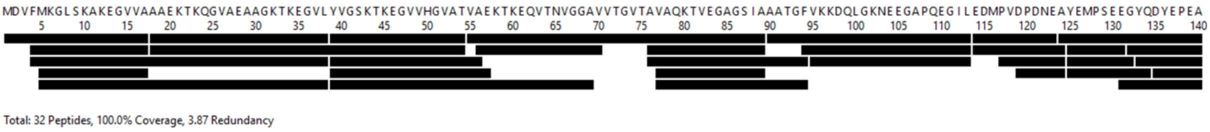


**Figure S5: Peptide coverage map in HDX-MS experiments. A total of 30 peptides were assigned with 100% aSyn sequence coverage and 3.87 degree of redundancy.**

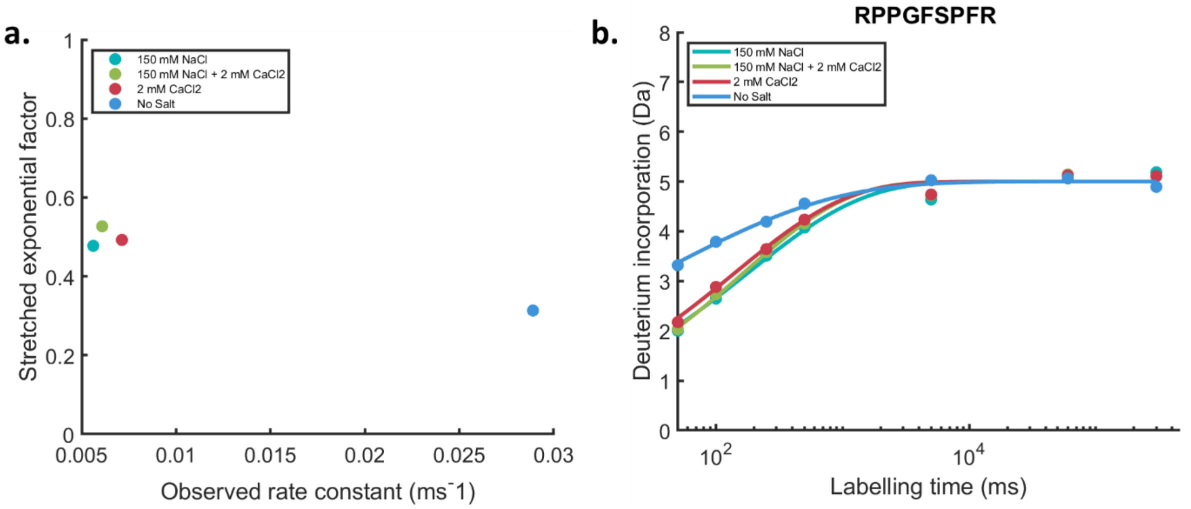


**Figure S6: Unstructured peptide bradykinin for the calibration of chemical exchange rate effects; a.** 2D plot of extracted fitted parameters $k_{obs}$ and β, with the “No Salt” condition (blue dot) chosen as reference state; **b.** Uptake curve for bradykinin in the four conditions; Data points are the mean of n=3 technical replicates.

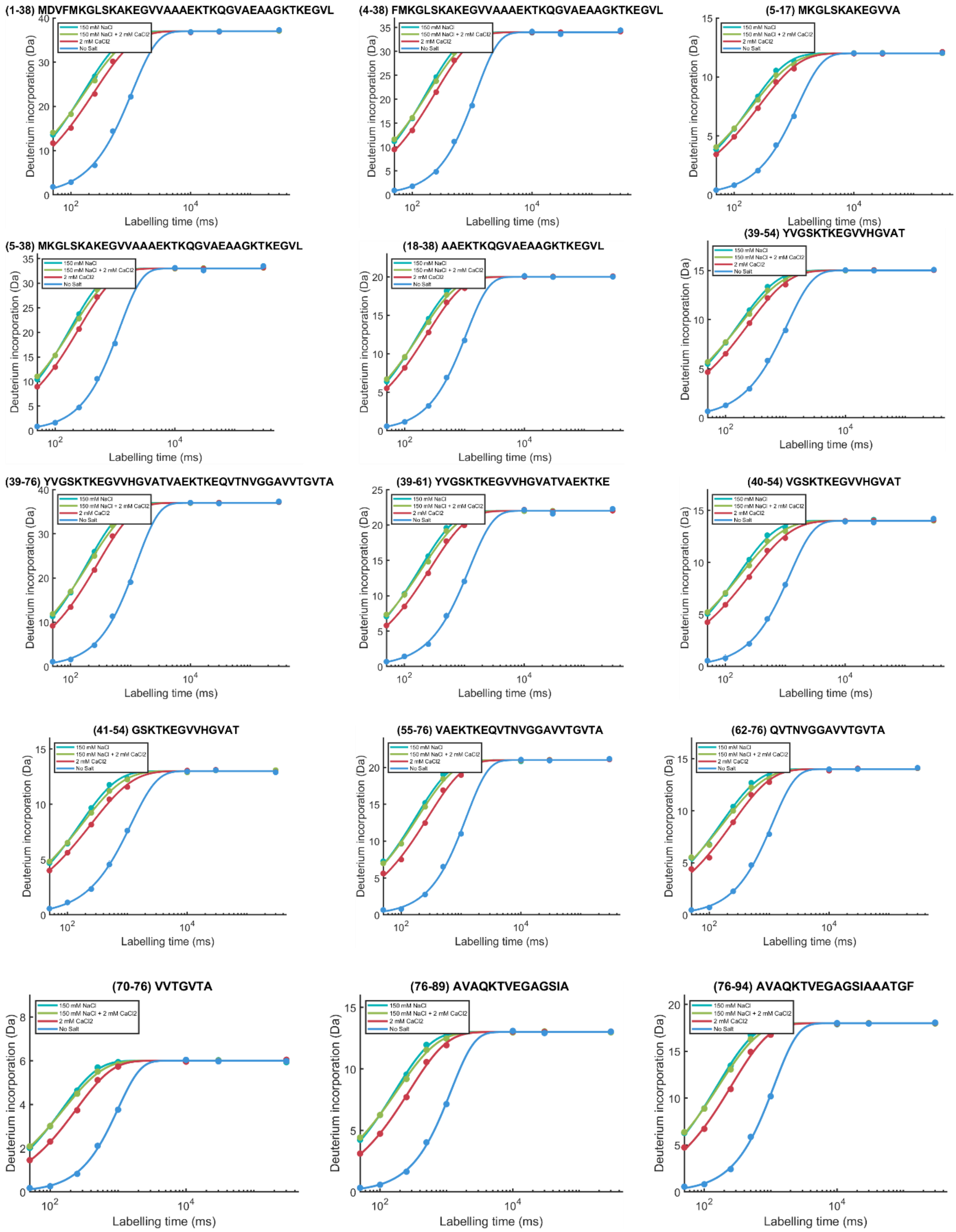

(1-38) MDVFMKGLSKAKEGVVAAAEKTKQGVAEAAGKTKEGVL
(4-38) FMKGLSKAKEGVVAAAEKTKQGVAEAAGKTKEGVL
(5-17) MKGLSKAKEGVVA
(5-38) MKGLSKAKEGVVAAAEKTKQGVAEAAGKTKEGVL
(18-38) AAEKTKQGVAEAAGKTKEGVL
(39-54) YVGSKTKEGVVHGVAT
(39-76) YVGSKTKEGVVHGVATVAEKTKEQVTNVGGAVVTGVTA
(39-61) YVGSKTKEGVVHGVATVAEKTKE
(40-54) VGSKTKEGVVHGVAT
(41-54) GSKTKEGVVHGVAT
(55-76) VAEKTKEQVTNVGGAVVTGVTA
(62-76) QVTNVGGAVVTGVTA
(70-76) VVTGVTA
(76-89) AVAQKTVEGAGSIA
(76-94) AVAQKTVEGAGSIAAATGF
150 mM NaCl
150 mM NaCl + 2 mM CaCl2
2 mM CaCl2
No Salt
Deuterium incorporation (Da)
Labelling time (ms)

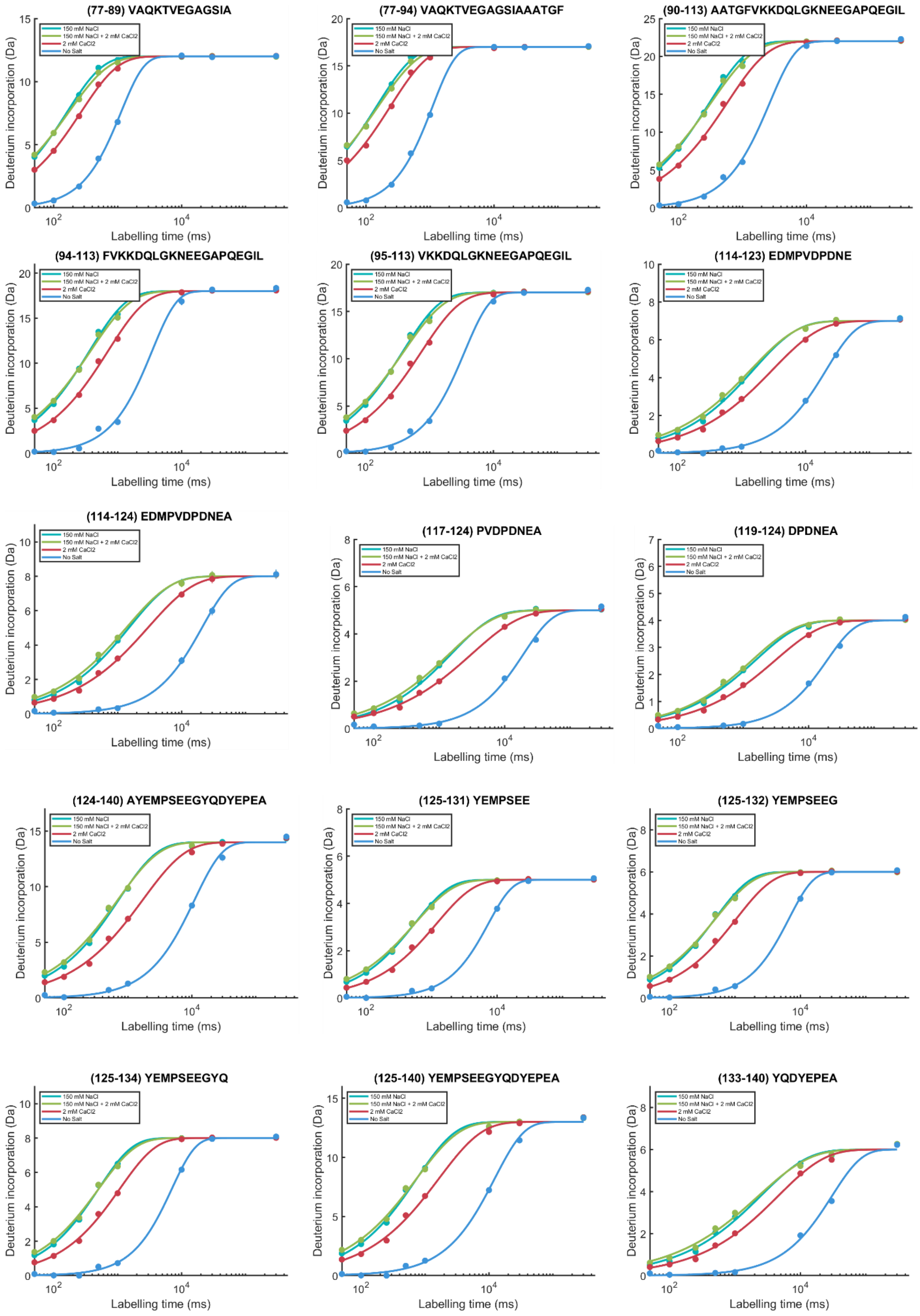


**Figure S7: Empirically adjusted deuterium uptake plots for aSyn equilibrated in No Salt, 2 mM $CaCl_2$, 150 mM NaCl, and 150 mM NaCl and 2mM $CaCl_2$ conditions.** Timepoints collected: 50 ms,

100 ms, 250 ms, 500 ms, 1000 ms, 10 s, 30 s, 300 s. Y-axis shows deuterium incorporation in Da and x-axis shows the log scale labelling times in ms. Data points are the mean of n=3 technical replicates.

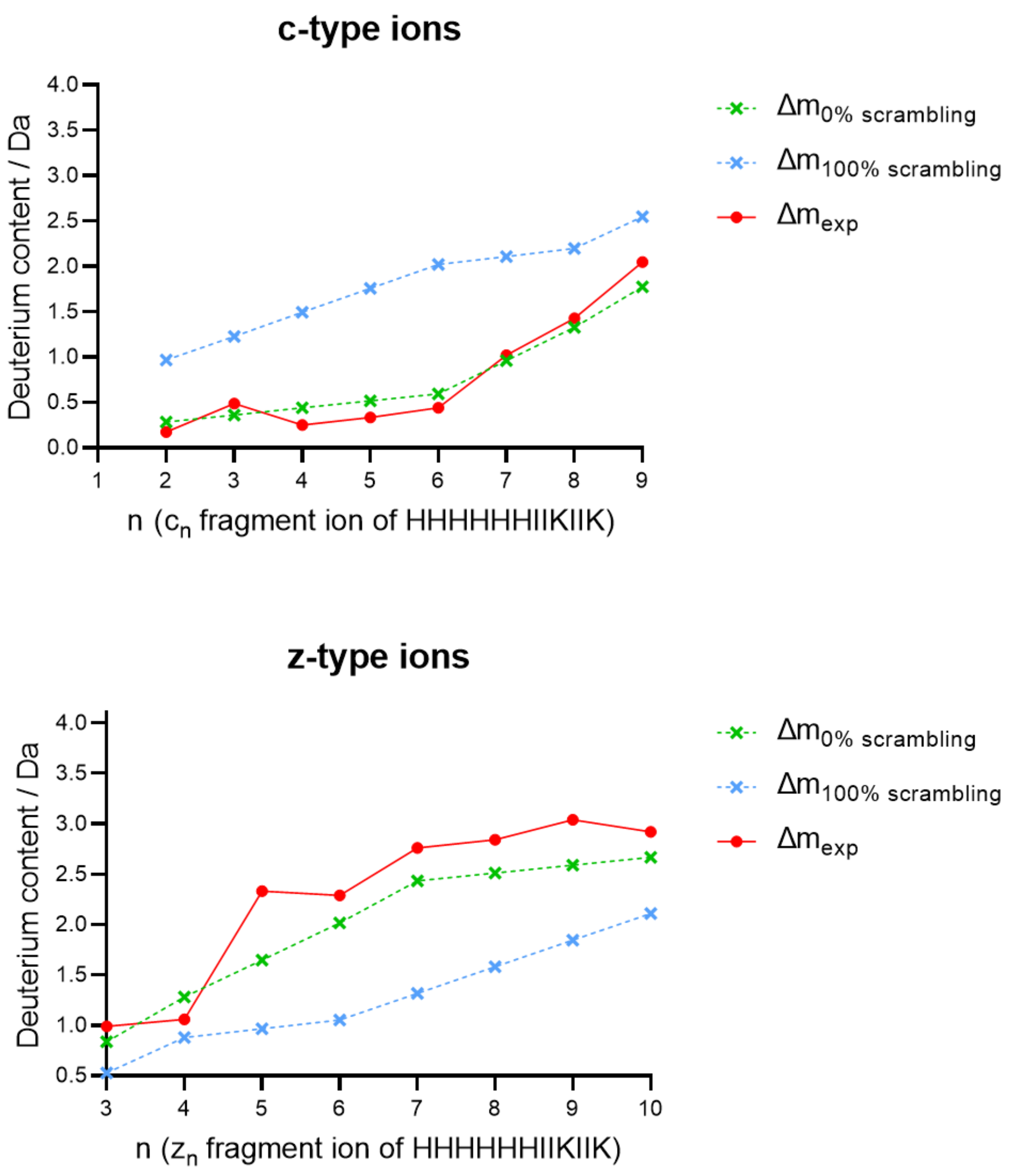


**Figure S8: Hydrogen-deuterium scrambling is not observed in the c and z fragments of peptide P1 under identical conditions as aSyn experiments.** The green and blue dotted lines on the plots represent the 0% and 100% theoretical scrambling data for this peptide respectively, and the red line shows the experimental data. From the plots, it can be inferred that the conditions chosen for ETD fragmentation do not cause H/D scrambling as the experimental data line (red) approaches the 0% scrambling line (green) for both types of fragments.

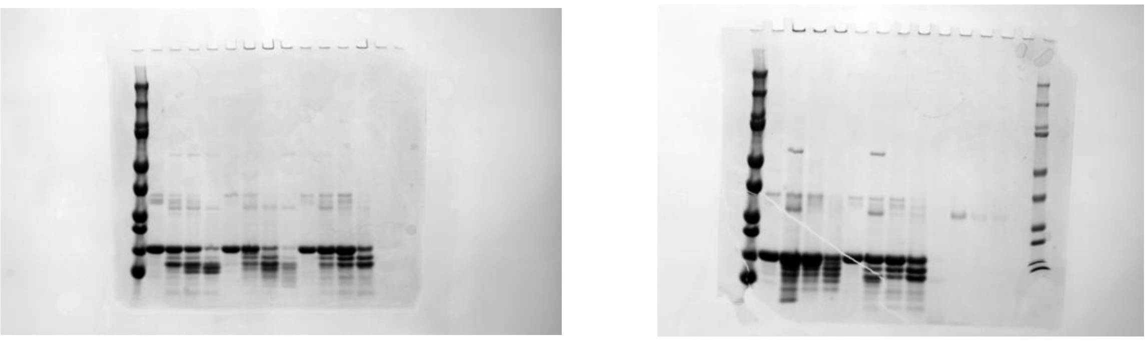

**Figure S9: Uncropped SDS-PAGE gels corresponding to proteinase K limited proteolysis experiments (Figure 6).**
